# Highly parallelized microfluidic droplet cultivation and prioritization on antibiotic producers from complex natural microbial communities

**DOI:** 10.1101/2019.12.18.877530

**Authors:** Lisa Mahler, Sarah Niehs, Karin Martin, Thomas Weber, Kirstin Scherlach, Miriam Agler-Rosenbaum, Christian Hertweck, Martin Roth

**Affiliations:** Bio Pilot Plant, Leibniz Institute for Natural Product Research and Infection Biology, Hans Knöll Institute, 07745 Jena, Germany; Faculty of Biology and Pharmacy, Friedrich Schiller University, 07743 Jena, Germany; Biomolecular Chemistry, Leibniz Institute for Natural Product Research and Infection Biology, Hans Knöll Institute, 07745 Jena, Germany; Physical Chemistry/ Microreaction Technology, Technical University Ilmenau, 98684 Ilmenau, Germany

## Abstract

To investigate the overwhelming part of the bacterial diversity still evading standard cultivation for its potential use in antibiotic synthesis, we have compiled a microscale-cultivation and screening system. We devised a strategy based on droplet-microfluidics taking advantage of the inherent miniaturization and high throughput. Single cells of natural samples were confined in 9 x 10^6^ aqueous droplets and subjected to long-term incubation under controlled conditions. Subsequent a high-throughput screening for antimicrobial natural products was implemented, employing a whole cell reporting system using the viability of reporter strains as a probe for antimicrobial activity. Due to the described microscale cultivation a novel subset of bacterial strains was made available for the following screening for antimicrobials. We demonstrate the merits of the in-droplet cultivation by comparing the cultivation outcome in microfluidic droplets and on conventional agar plates for a bacterial community derived from soil by 16S rRNA gene amplicon sequencing. In-droplet cultivation resulted in a significantly higher bacterial diversity without the common overrepresentation of Firmicutes. Natural strains able to inhibit either a Gram-positive or a Gram-negative reporter strain were isolated from the microscale system and further cultivated. Thereby a variety of rare isolates was obtained. The natural products with antimicrobial activity were elucidated for the most promising candidate. Our method combines a new cultivation approach with a high-throughput search for antibiotic producers to increase the chances of finding new lead substances.

## Introduction

In light of the antibiotic resistance crisis^1^, there is a dire need for fundamentally new lead structures which can be altered into antibiotics. The microbial diversity provides a rich resource for new chemical scaffolds^2^ and fortunately, the microbial diversity is far from being exhaustively mapped. On the contrary, it seems like after one and a half century of microbiological research we have only started to understand the true extent of the microbial variety, initiated by culture-independent methods like sequencing of rRNA cloning libraries in the late 1980s^3^. The fact that only 1% to 15% of the bacterial diversity are cultivable under standard laboratory conditions^4^ promises further revolutionizing compounds to be discovered in the remaining majority. The question is how we can unlock the hidden biosynthetic potential of previously uncultured microbes. Axenic cultures are mandatory to study the regulation as well as the chemical and biological characteristics of natural products and hence, appropriate conditions are needed for cell replication of species recalcitrant to cultivation.

To narrow the gap between the naturally occurring complexity of microbial communities and the limited diversity of culture collections, several new techniques have been developed. For instance, concentration and composition of media have been adapted for the large number of oligotrophic uncultured species^5–10^. Other techniques mimic the conditions *in situ* in more detail by supplying extracts^5,11–13^ derived from the environment containing unspecified mixtures of macro- and micronutrients and other growth factors. The most advanced development in this direction is to provide direct contact with the environment^14–17^, which proved highly productive in culturing species regarded as unculturable.

A complementary trend is the miniaturization of the culture vessels^12,14,18–30^ to save material and time while increasing the throughput and thereby the chance to find non-dormant, culturable variants of naturally occurring species. These microscale culture techniques have the idea in common to partition complex bacterial communities on the single cell level, allowing the various species to grow at their own speed, since they do not have to compete neither for space nor for nutrients. At the same time, the compartments that confine single cells are small and densely packed to establish a highly parallelized cultivation. The reported microscale approaches can be divided into methods that arrange the compartments into arrays^18,20,23–30^ and techniques that keep the compartments not fixed to positions but rather mobile within the population^12,14,19,21,22^. The clear advantage of arrays is the possibility to address the same culture several times, since the compartments are identified by their coordinates. Ingham *et al*.^24^, for instance, created an exceptional array called the micro-Petri dish with 180,000 wells of 20 μm x 20 μm and used 25 of them to screen a Rhine water community for the ability to dephosphorylate a fluorescent substrate. Positive colonies were picked by hand.

However, array technologies also have some disadvantages. First, space is lost by introducing a distance between array positions for further downstream manipulation and by omitting the expansion in the 3^rd^ dimension, since up to now all arrays are only two-dimensional. Second, fast automation is simpler with moving compartments because slow high precision positioning systems or manual selection processes to interface the arrays are not necessary but only methods to manipulate the direction of the moving vessels^31^. Consequently, higher absolute numbers of compartments can be generated and processed per unit of time. For instance, Zengler *et al*.^12^ generated 10,000,000 droplets in bulk inoculated with aquatic or soil communities and screened for growth by side scatter measurement in fluorescence activated cell sorter (FACS) with 5000 droplets per second.

We propose a cultivation and screening strategy based on surfactant-stabilized microfluidic droplets in a perfluorinated oil phase. Microfluidic droplets provide a genotype-phenotype linkage also for secreted products, meaning that those compounds stay confined with the cell from which they originated. The droplets have been stored in bulk, allowing to generate and incubate populations of 9 x 10^6^ droplets under controlled aerobic conditions. Furthermore, we have implemented a high throughput screening for antimicrobial products using whole cells as reporting system in droplets^32^. We verified our cultivation and screening strategy by culturing a complex bacterial community derived from a soil sample in a growth substrate containing low concentrations of cold extracted soil extract, soil particles and plant derived nutrients. Bacteria selected from these cultures due to their bioactivity against *Escherichia coli* were identified as candidates for a new genus by sequencing.

## Results

### Concept of cultivation in pL-droplets and workflow

We used a droplet-microfluidic platform to singularize, cultivate and screen environmental cells from complex bacterial communities. The workflow featured three major phases: First, cells were separated from each other by encapsulating them in droplets during droplet generation. All droplets were collected in a vessel, where they lost their order. We refer to the state of a droplet population, in which the positional information of the droplets is unrelated to the order of droplet generation, as “droplets in bulk”. Second, droplets in bulk were incubated together using dynamic droplet incubation^33^, which ensured aerobic and homogenous cultivation conditions for weeks or months. Third, grown microbial cultures were isolated from droplets by depositing them on agar plates. Either all droplets were deposited in an untargeted manner or droplets were first screened for antibiotics and then, only selected droplets were distributed on agar plates.

During droplet generation, we co-encapsulated low amounts of nutrients originating from plants, a cold extracted soil extract (CESE) prepared from the same soil that the community was derived from, and several solid soil particles (diameter < 40 μm). While designing the medium we followed the rationale of replicating the conditions in the natural environment as closely as possible. The number of cells occupying a droplet at encapsulation follows a Poisson distribution^34^. As we aimed at one cell per droplet, we minimized the probability of several cells per droplet by adjusting the concentration of cells upon inoculation to obtain not more than 30% occupied droplets (λ= 0.4, in average 0.4 cells/droplet). Nevertheless, the occurrence of several cells per droplet cannot be excluded due to a small statistical probability of 6.16%, and biological reasons like cells adhering to the same soil particle or to each other. The droplets were stabilized by a biocompatible surfactant forming a monolayer at the aqueous/oil interphase. Surfactant addition reliably prevents coalescence of droplets upon mere droplet-droplet contact or cell cross-contamination over several droplets, thereby allowing to incubate several millions of droplets together in the dynamic droplet incubation setup. The level of miniaturization (~200 pL droplet volume) in conjunction with bulk incubation, permitted droplet populations comprising on average 9 x 10^6^ droplets.

### In-droplet cultivation results in higher CFU concentration in comparison to conventional plating

We used a brown earth soil sample to extract an environmental microbial community as well as a cell-free extract, termed CESE, containing dissolvable macro- and micronutrients and other effectors, like signaling molecules. To prove the capability of our droplet cultivation technique, we directly compared the cultivation outcome of droplets with that of standard plating. The soil community was incubated with the same media composition at 20 °C for 28 days in both approaches. We compared one droplet population of approximately 9 x 10^6^ droplets to 15 agar plates, including 8 plates inoculated with a 1:100 and 7 plates with a 1:1000 dilution of the original community. This comparison was replicated 8 times.

During incubation we monitored microbial growth by brightfield microscopy imaging of the droplets followed by counting occupied droplets (Figure 1). After 28 days the average occupation rate of the 8 populations amounted to 22.4%, which translates into 3.03 x 10^6^ colony forming units (CFU)/mL of the soil community preparation at an average droplet volume of 200 pL (Figure S1). At the same time, we counted 387 colonies per plate on average for the 1:100 dilution and 92 colonies per plate for the 1:1000 dilution. On plates, the colony count is equivalent to 1.3 x 10^6^ CFU/mL of the soil community (0.77 x 10^6^ CFU/mL for 1:100 dilution and 1.83 x 10^6^ CFU/mL for 1:1000 dilution respectively). The more than two-fold higher concentration of CFUs found during in-droplet cultivation indicates that singularization and incubation of cells in droplets enabled more cells to replicate. We suggest this is not only due to a more sensitive detection of cells in droplets, aided by microscopy, but also to the growth of cells that would not form colonies on agar plates.

**Figure 1.**
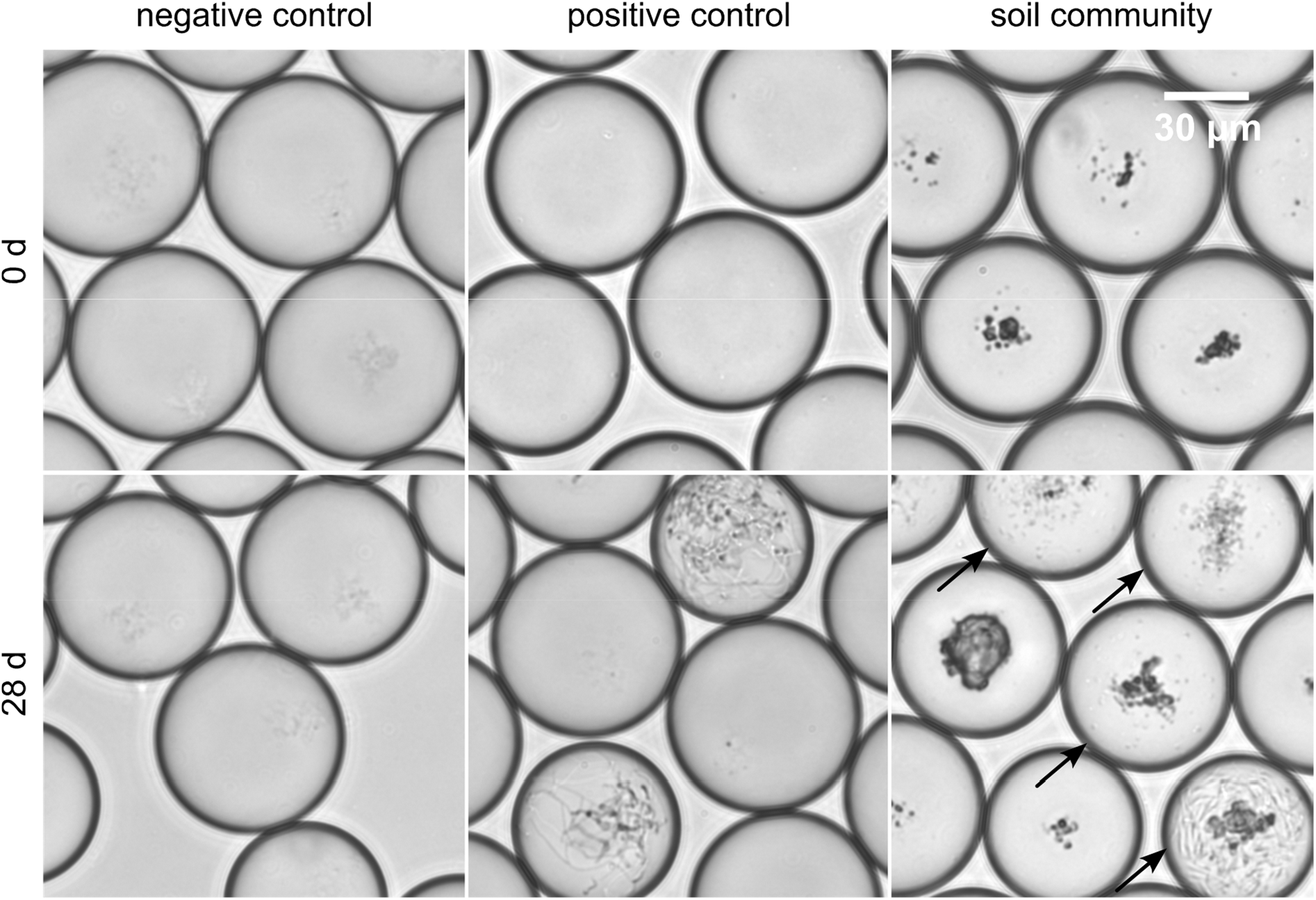
Brightfield microscopy images of droplet samples after droplet generation and after 28 d of incubation. Images were taken on an inverse microscope at 40 x magnification while droplets were trapped in an observation chamber. Every 7 days, ca. 5 μL of droplets were sampled from the dynamic droplet incubation setup to examine microbial growth. Bacterial colonies were not visible right after droplet generation (top). Negative control droplets remained empty throughout the entire incubation period (bottom left). Co-encapsulated soil particles are visible as dark particles in the images of droplets with soil community (right). Accumulation of bacterial cells was already detectable after 7 days of incubation in the positive sample and in droplets inoculated with soil community (not shown). A variety of different morphologies was observed in droplets with soil community, ranging from different sizes and densities of cocci and rod shaped cells to filamentous growth (bottom right). Droplets with cell colonies are marked with arrows.

### A diverse set of microbial species replicates in droplets

We analyzed the diversity of growing cells in 8 replicates for each cultivation method by pooling the biomass of the colonies after incubation, extracting metagenomic DNA and sequencing the 16S amplicons on an Illumina MiSeq platform. For comparative analysis we subsampled the reads of all samples to an equal depth of the lowest number of reads (88435 in sample D_SM8) to avoid bias by variable library size^35^. In order to reduce noise in the data set, uninformative OTUs were removed by restricting the dataset to OTUs represented by at least three reads in at least three samples. The combination of a total count and prevalence filter reduced the number of OTUs from 330,329 to 2,106. Furthermore, a median normalization within samples was applied to the read counts.

To explore the similarity between the microbial community structures for the different cultivation approaches, we performed unconstrained ordination based on the community composition using the Bray-Curtis distance (Figure 2 A). The spread of the data correlated well with the sample type (e.g. samples from droplet cultivation vs. samples from plate cultivation), meaning that droplet samples clustered closely without overlapping with plate samples. Samples belonging to one sample type showed little dispersion. These results indicate clear dissimilarities between the cultivation outcome of the two methods but a reproducible community composition within one cultivation method.

**Figure 2.**
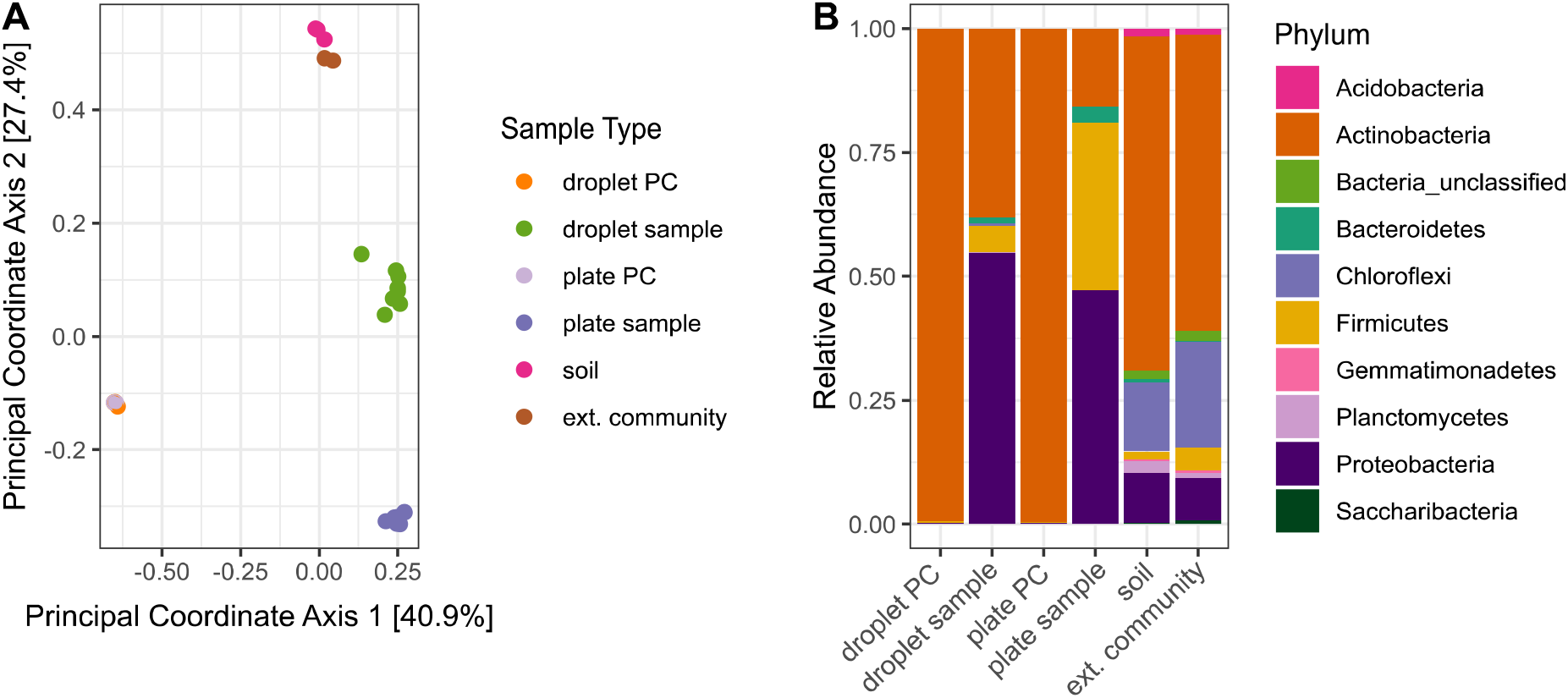
Differences in bacterial diversity due to cultivation technique. Sample type abbreviations denote: droplet PC – droplet positive control, droplet sample – droplet population inoculated with soil community, plate PC – plate positive control, plate sample – agar plates inoculated with soil community, soil – original soil sample, ext. community – extracted soil community. A) Principal Coordinate Analysis based on Bray-Curtis distance matrix visualized similarities between microbial communities. Abundances of OTUs (evolutionary distance of 0.03) with a minimum of 3 reads in at least 3 samples were normalized to the corresponding sample median and used for the ordination. B) Community structure based on the Illumina amplicon sequencing of the 16S rRNA gene. The phylogenetic affiliation at the phylum level is displayed for each sample type. The bars depict the relative abundance of the 10 most abundant phyla.

At the broad level of phyla, differences in the community structure are clearly visible between all sample types (Figure 2B). The relative abundance of the phylum Actinobacteria among the cultivated species was more than two-fold higher in droplets (37.9%) than on agar plates (15.7%). As expected more than 99% of the reads in the positive controls for both cultivation techniques affiliated to Actinobacteria, implying no contamination of the cultivation approaches and an appropriate taxonomic classification. The composition of the original soil sample was similar to the initial community structure of the extracted soil community.

Alpha diversity indices were calculated for the subsampled dataset, but no further filtering was conducted. The total number of OTUs was significantly higher in the droplet cultivation samples than in the plate cultivation samples, as was the number of Singletons and, consequently, the Good’s Coverage index (Figure S2). The higher number of Singletons indicates that even with a sequence depth of almost 90.000 reads, the probability to have missed part of the diversity is higher than in the agar plate samples, also implying a higher degree of diversity in droplets. Significantly higher chao1 and Shannon indices in droplet samples, as estimates for species richness and species diversity, respectively, further supported a higher microbial diversity cultivated in droplets (Figure S2).

To examine the differences in diversity in more detail, we directly compared the relative abundance of taxa on all taxonomic ranks for the cultivation outcome of droplets and plates (Figure S4-S8). For both cultivation techniques, we found as most abundant phylum the very diverse Proteobacteria (droplets 54.7%, plates 47.3%), followed by Actinobacteria (31.6%) for droplets, and Firmicutes (33.7%) and Actinobacteria (14.7%) for plates. While on plates the majority of Proteobacteria was constituted by Alphaproteobacteria (90.8%), the phylum split up almost equally into Alpha- and Gammaproteobacteria (57.3% and 37.7%) in droplets. Another pronounced distinction was the significantly higher abundance of the phylum Firmicutes on plates, comprising one third of the community (Figure S4), while it was represented with only 5.3% in droplets. On plates the Firmicutes were mainly composed of the genus *Bacillus* (55%) and *Paenibacillus* (18%) to which 26 and 25 OTUs affiliated, respectively. However, both genera were strongly biased to only 3 (*Bacillus*) or 2 OTUs (*Paenibacillus*). As expected, *Bacillus* and *Paenibacillus*, together with *Phyllobacterium*, were the three most abundant genera on plates, accounting for 42% of the sequences (Figure 3 A). In contrast, *Acinetobacter* as member of the γ-Proteobacteria showed the highest relative abundance on the genus rank for droplets, accompanied by *Agromyces* and *Micromonospora*. Together they covered 44% of the reads. The two latter genera both belong to the phylum Actinobacteria, whereas *Micromonospora* are a rich source of antibiotic compounds, like aminoglycosides. Also, other genera assigned to Actinobacteria, like *Solirubrobacter, Nocardioides* and several uncultured members of the family Acidimicrobiaceae, were found to be significantly more abundant in droplet samples. Remarkably, the well-known genus *Streptomyces* was observed more often on plates. As a general trend, most taxa present on plates were also found in droplets, while many taxa were exclusively found in droplets but not on plates (Figure 3 B, C, S3 A). Both cultivation methods enriched a few species in high abundance, which dominated almost 50% of the sequences, but in droplets the remaining 50% were more evenly comprised of a richer set of microorganisms. Besides the taxa that were exclusively found in droplet samples, accounting for almost 60% of all taxa, we found also a considerable number of taxa classified as uncultured on the genus level among the exclusive droplet taxa (151 taxa of 992 exclusive droplet taxa) (Figure SI 3 B, C). Taxa that were just unclassified and likely belong into the uncultured category were not considered for this comparison. Among the 151 uncultured droplet taxa were 7 taxa belonging to the phylum Saccharibacteria, which in turn belongs to a large cluster of lineages mostly lacking cultured representatives^36^. While 15.2% of the exclusive droplet taxa were classified as uncultured only 1.2% of the taxa found in both, droplet and plate samples, were classified as uncultured and 0% in exclusive plate taxa.

**Figure 3.**
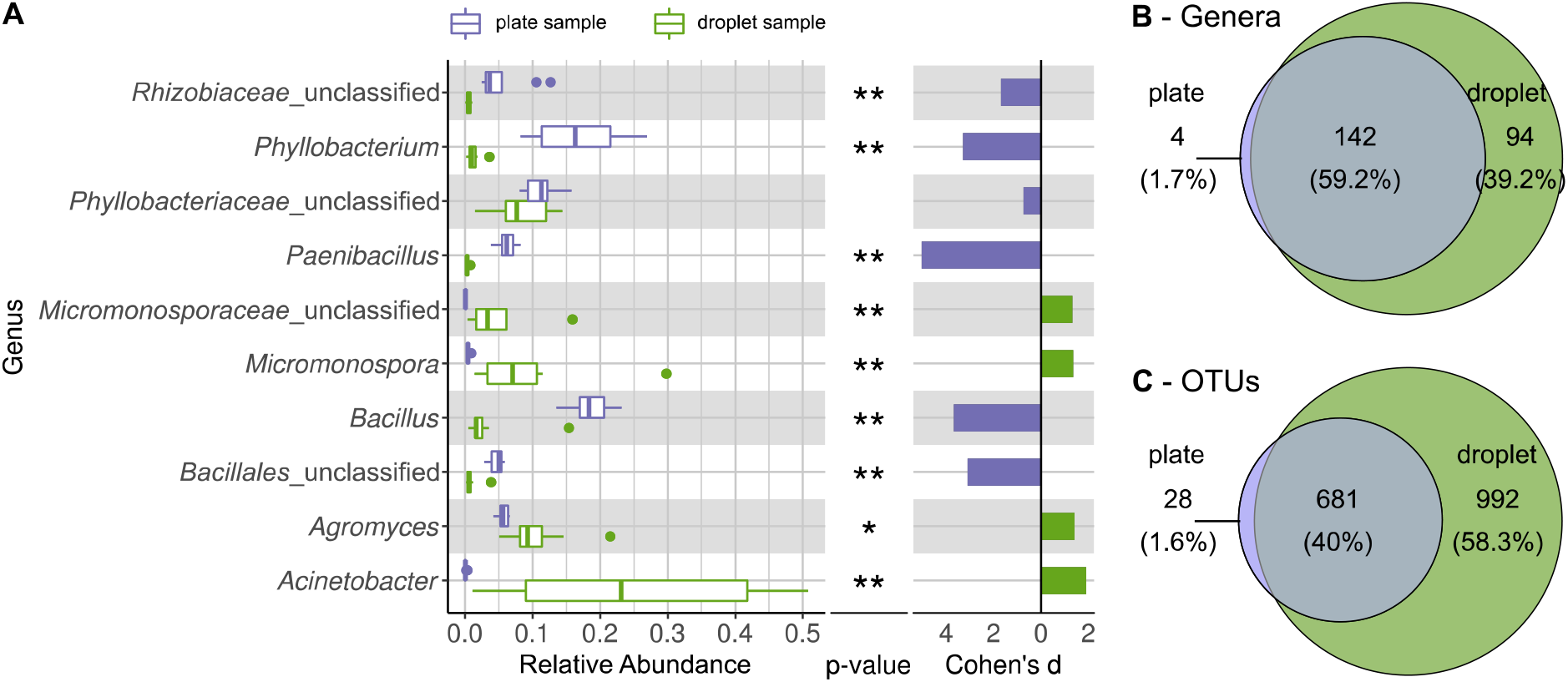
Comparing taxonomic classification on genus rank for replicating cells in agar plate and in-droplet cultivation. A) Boxplots depict the distribution of relative abundances for genera in droplet and plate cultivation samples. Relative abundances of OTUs (evolutionary distance of 0.03) were agglomerated on the genus level (evolutionary distance of 0.1). Displayed are the 10 most abundant genera of in total 240 assigned genera covering 66.8% of all sequences. Means of relative abundances were compared by Wilcoxon Rank-Sum test for each genus (α=0.05), applying Holm-Bonferroni correction for multiple comparisons. ** significant with *p* < 0.01, * significant with *p* < 0.05. As effect size Cohen’s *d* was computed and plotted as bars to indicate which differences are practically relevant. The direction and the color of the bars depend on the sample type in which the larger mean was found (blue – larger mean in plate samples, green – larger mean in droplet samples). B) Venn diagram for the taxa on genus level. Of 240 assigned genera 142 were found in both cultivation methods while 94 were unique to droplet cultivation and 4 were only observed in plate cultivation. The area of the Venn elements correspond to the total number of genera found for the cultivation techniques. C) Venn diagram for the OTU level. Of 1701 OTUs, 681 were found in both cultivation methods, 992 were unique to droplets and 28 were unique to plates. The area of the Venn elements correspond to the total number of OTUs found for the cultivation techniques.

### Microbial colonies can be isolated from microfluidic droplets

To facilitate the isolation of bacterial cells from droplets, we designed an experimental setup, in which droplets were deposited on a nutrient containing matrix after one month incubation in order to enable microcultures to grow into macroscopically visible colonies (Figure 4). Via a capillary, into which droplets were injected at 10-50 droplets/s, droplets were guided to an agar plate on a positioning system. While droplets and oil phase continuously emanated from the capillary tip, the plate moved following a spiral pattern, leading to the distribution of droplets along the spiral over the agar plate. Both the volatile oil phase surrounding the droplets and, subsequently, also the aqueous droplets evaporated fast, leaving the cells behind that were previously confined in the droplets now spatially separated on the agar surface. After incubation of the agar plates, macroscopic colonies were picked, purified by restreaking and characterized.

**Figure 4.**
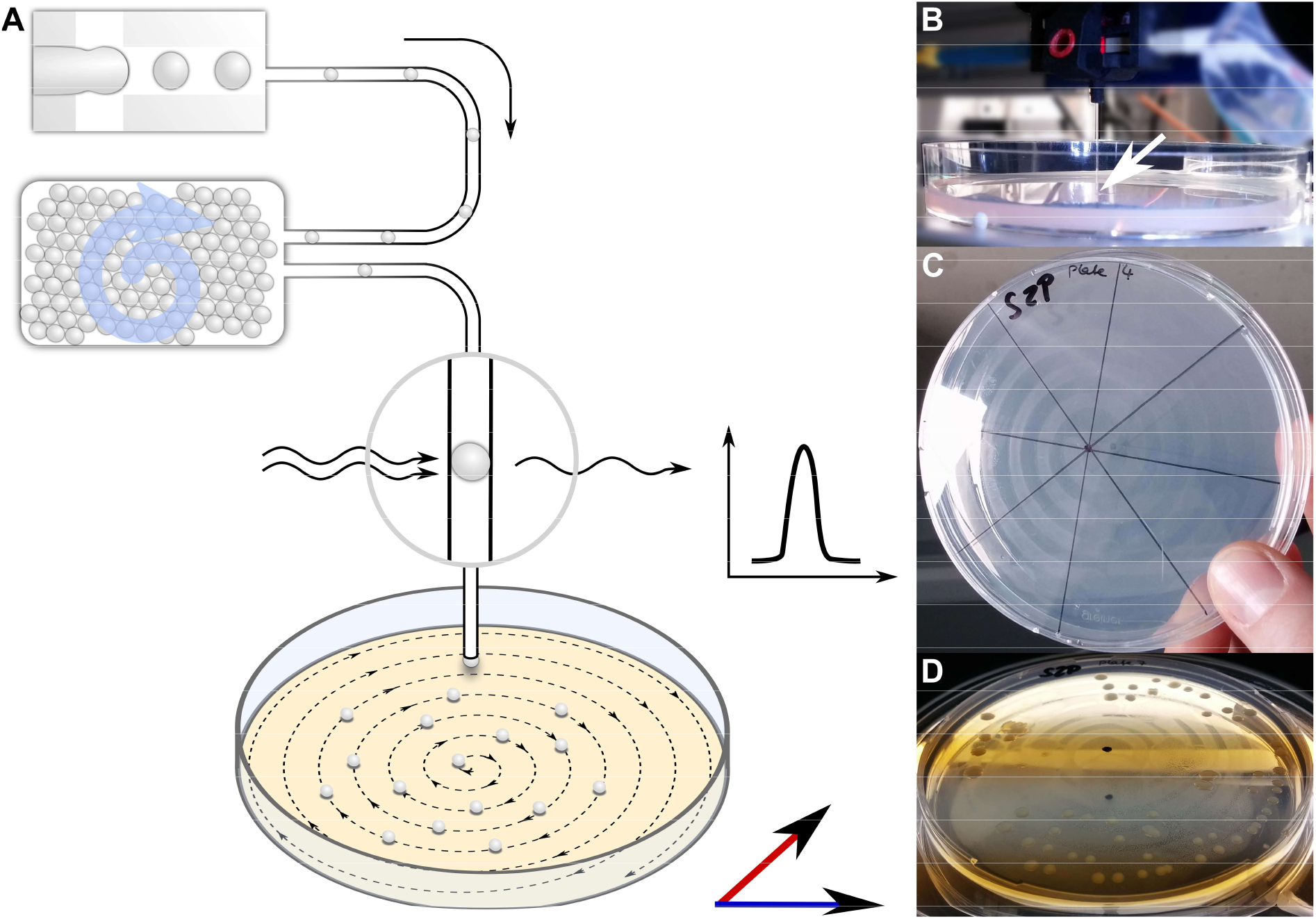
Continuous droplet deposition on an agar plate of preincubated droplets. A) Sketch of continuous droplet deposition workflow. Droplets inoculated with cells of the soil community were incubated for one month, before they were injected into a capillary (100 μm ID) positioned above an agar plate. The agar plate moved in x-y direction at a speed of 1 mm/s, while the droplets were injected at 10-50 droplets/s. The droplet frequency was monitored by a custom-developed sensor. B) Image of the capillary taken while droplets are deposited on an agar plate. The tip of the capillary is highlighted by a white arrow. The capillary tip is close to the agar surface, in order to establish a continuous fluid film. C) An agar plate immediately after droplet deposition with the dried oil film in the spiral pattern of continuous deposition. D) An agar plate with deposited droplets after incubation for two weeks. The dried oil film is still visible, on which the bacterial colonies are now aligned.

We picked 224 colonies in total, trying to keep the selection random and thereby reflecting the true distribution of abundance for the bacterial diversity. 301 axenic isolates were obtained on agar plates, of which 266 were initially characterized by sequencing the full 16S rRNA gene. The isolates covered four major phyla (Firmicutes, Actinobateria, Proteobacteria, Bacteroidetes) resolving in 11 different genera (Figure 5). Most of the isolates were classified as *Bacillus* (52.3%), underlining the strong bias towards Firmicutes caused by intermediate cultivation on agar plates. Nevertheless, the accumulated abundance of Actinobacteria amounted to 30.5%, resembling the abundance in the amplicon data. Moreover, the Actinobacteria were represented by several rarely observed genera like *Kocuria* and *Leifsonia*, and a notable high frequency of *Micromonospora* with 22.9%, which showcases the influence of in-droplet cultivation. Remarkably, one of the isolates clustered only to reference sequences of uncultured bacteria assigned to the family Cytophagaceae. However, this isolate could not be maintained for long on standard agar plates. The obtained genomic DNA could nevertheless be further investigated.

**Figure 5.**
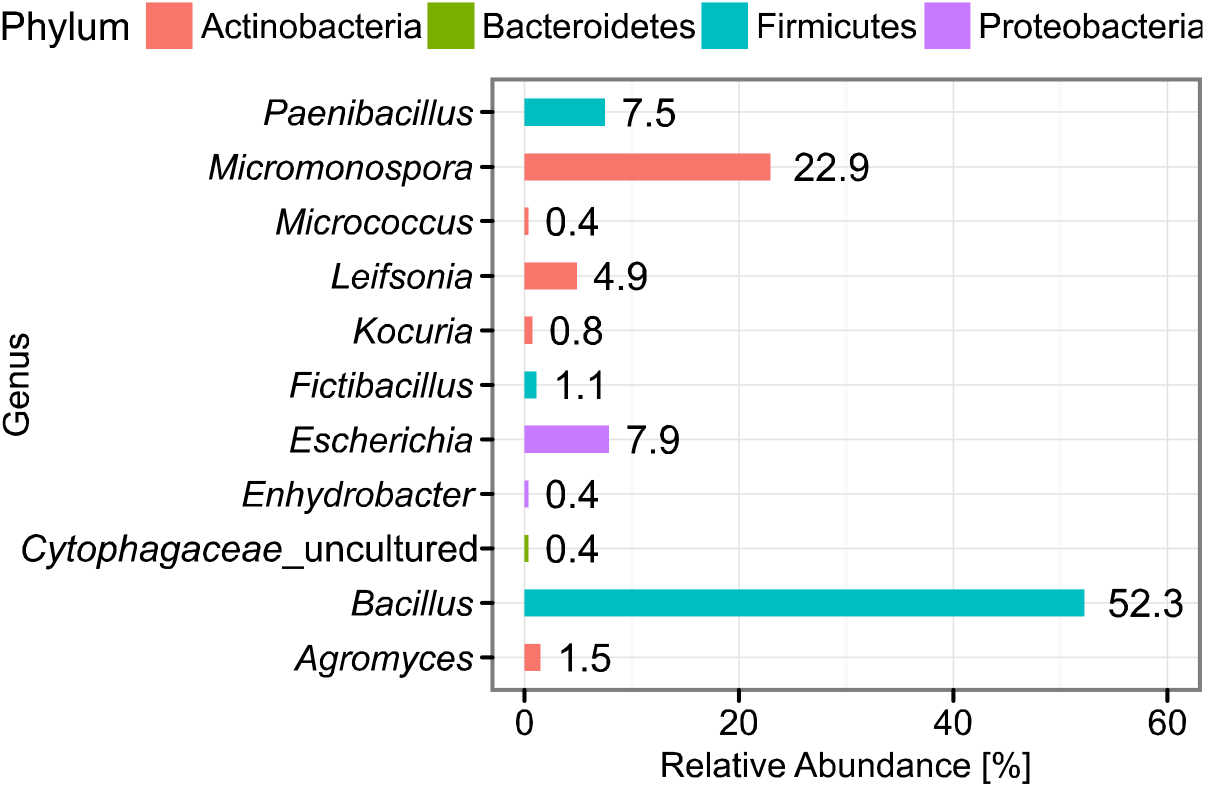
Abundance of classified bacterial isolates obtained from droplet cultivation. The distribution of abundance on the genus rank is displayed. Colors of the bars indicate the affiliation to phyla. Each bar is labeled with its exact relative abundance.

A bias towards isolates compatible with agar plate conditions and growing to high cell densities is unavoidable here and optimization is in progress, but we hypothesize a droplet aided strain discovery is anyway possible using this method. We base our hypothesis on a combination of the scout model, postulating that cells of a dormant population switch stochastically into an active state scouting the current conditions, and spontaneous domestication as described by Buerger and Epstein^37–39^. The high parallelization in droplet cultivation leads to substantially increased chances to find rare scout cells, which can replicate and by doing so form even more rare culturable variants, termed spontaneous domestication. Those variants would grow on agar plates and are more likely to be found with the droplet workflow.

### Screening for antibiotic producers in droplets

Microfluidic droplets are extraordinarily suited for screening applications since a fast and automated read-out is inherent to the system. We combined the cultivation of complex cell mixtures with an immediate screening for antibiotic compounds^32^ in order to achieve the selective isolation of putative producers of antimicrobial substances. For this purpose, we pico-injected cells of a reporter strain into all droplets of a population after one month incubation (Video S1). The reporter strain expressed a gene for a red fluorescent protein (RFP) as a survival signal in case its metabolism was functioning normal. In the presence of an inhibiting compound produced by the cultivated environmental cells, replication of the reporter and expression of RFP were disturbed resulting in a reduced fluorescence intensity. After the environmental cells had been co-incubated with the reporter cells inside the droplets for one additional day, we used the fluorescence intensity of the reporter to classify droplets into two categories, based on which we sorted the droplets in a microfluidic sorting chip: i) Droplets with inhibited reporter strain showing a low fluorescence signal and ii) droplets with uninhibited reporter strain showing a high fluorescence signal. Droplets with low red fluorescence signal were selected in a droplet sorting operation (Video S2) and subsequently distributed on agar plates to recover soil bacteria producing antimicrobial compounds.

In four different screening experiments, we explored combinations of two different reporter strains, *Escherichia coli* and *Bacillus subtilis* as representatives of Gram-positive and Gram-negative bacteria, and two different media to cultivate the soil community in droplets (Table S2). The recovered isolates of the selected droplets were characterized by 16S rRNA sequencing. We observed an enrichment of Actinobacteria and Alphaproteobacteria represented by various genera like *Kocuria, Microbacterium* and *Sphingomonas* for the *B. subtilis* reporter (Figure 6, Figure S9). In contrast, the spectrum of genera recovered with the *E. coli* reporter was broader. The most abundant genus was however *B. subtilis*, which was also the case for the *B. subtilis* reporter. Interestingly, two isolates falling in the Cytophagaceae family but being only distantly related to known type strains (<95%) were obtained in the *B. subtilis* screen. Also, the Cytophagaceae isolate gained in the untargeted isolation did not cluster closely (Figure S10), indicating that those isolates represent an additional new strain. As before, these isolates could not be maintained on agar plates for long.

**Figure 6.**
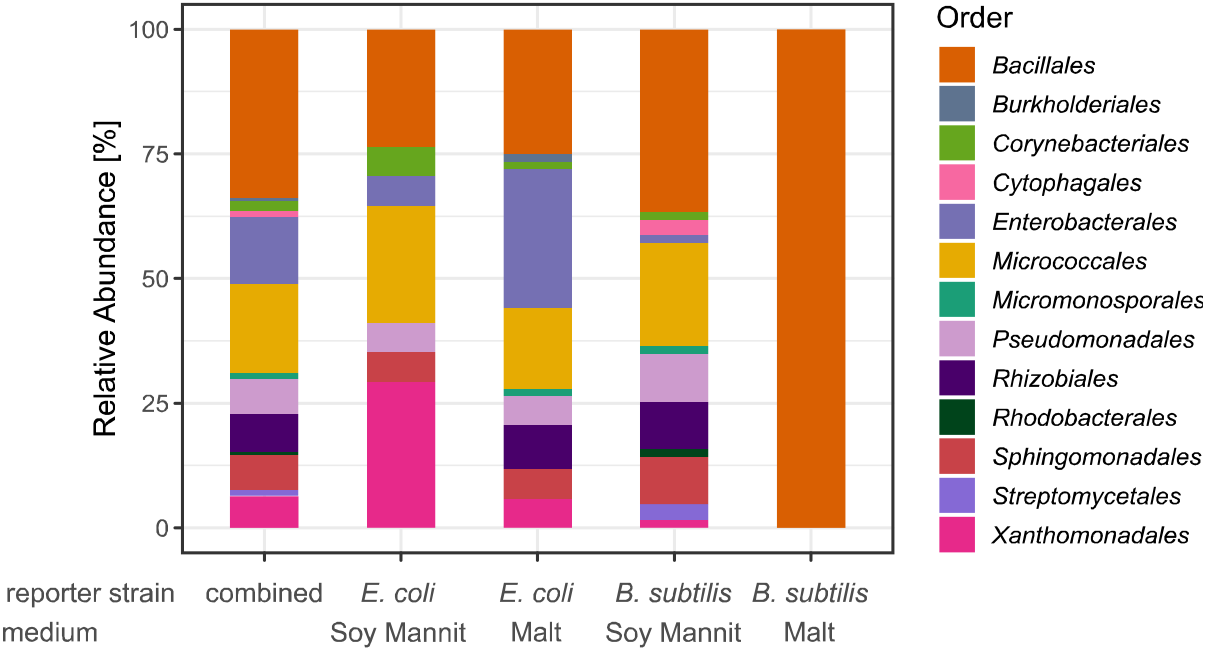
Taxonomic diversity of isolates obtained in the screening for antibiotic producers. The classification is depicted on the level of order and based on the Sanger sequences of the nearly complete 16S rRNA gene. The bars depict the relative abundance of all detected orders for the isolates of all 4 screening rounds combined and for each screening round separately.

### Natural product reservoir

We isolated a closely related strain of *Bacillus tequilensis* (99.79%; isolate D121-0906-b3-2-1) that showed strong growth inhibitory properties towards various bacterial strains like *Mycobacterium vaccae* or *Pseudomonas aeruginosa* during the primary validation of our screening hits. Additionally, the crude culture supernatant exhibited high antifungal activity against *Candida albicans* and *Penicillium notatum*. For increasing the production titer, we cultured *Bacillus* strain D121-0906-b3-2-1 under several cultivation conditions including varying media composition, duration of cultivation, and extraction method. Based on exact masses and the characteristic UV absorption spectra, we were able to detect bacillaene A and B, strong antibacterial polyketides active against a variety of strains^40^ (Figure 7, S11). In addition, highly lipophilic metabolites could be detected with exact masses ranging from 993–1053 Da. Characteristic amino acid fragments obtained using HRESI-MS/MS revealed these cryptic molecules to be gageostatins A–C^41^, cytotoxic lipopeptides that are known to exert good antifungal activities (Figure 7, S12-S14).

**Figure 7.**
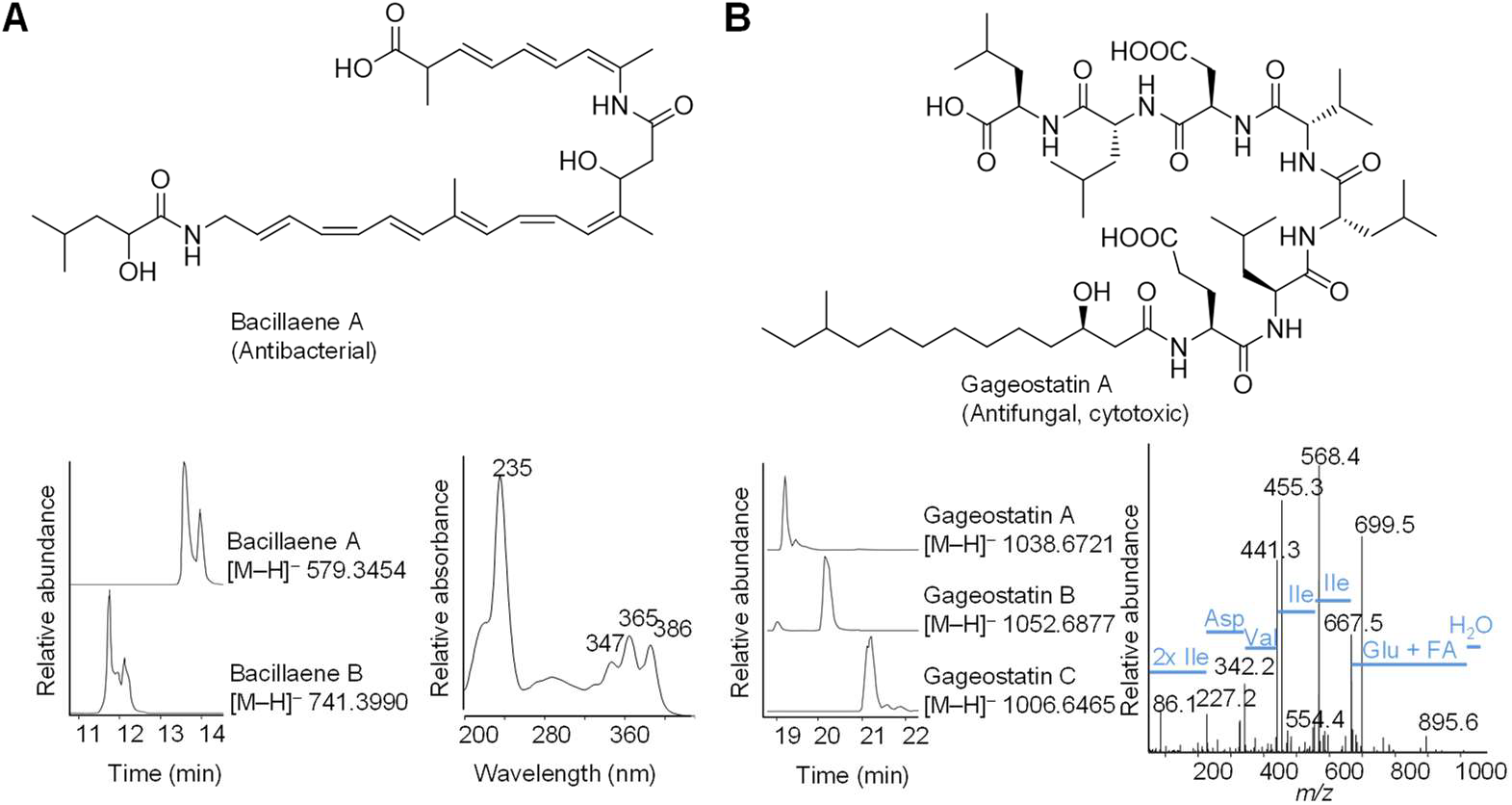
Detected natural product arsenal of *Bacillus* sp. strain D121-0906-b3-2-1. A) Production of bacillaenes in metabolic profile of *Bacillus* sp. strain D121-0906-b3-2-1 extracts and characteristic UV absorption spectrum for bacillaene A. B) Production of gageostatins and MS/MS fragmentation pattern for gageostatin A. FA – Fatty acid. All LC/MS profiles are displayed as extracted ion chromatograms in negative mode.

## Discussion

We devised an innovative microscale cultivation strategy utilizing a modular droplet-microfluidic platform and mimicking environmental conditions *in situ*. The miniaturized droplet volume of 200 pL facilitated high density cultivation of 2 million compartmentalized single cells in a total volume of 1.5 mL (9 million droplets with 22% occupancy on average). By choosing to conduct cultivation in surfactant-stabilized aqueous droplets permanently surrounded by oil, we achieved robust confinement of cells over long periods of time. Additionally, we were able to screen for antibiotic production, which demanded the addition of reporter strains to the cultivation compartments.

One major advantage of our method compared to previous microscale cultivation systems lies in the space efficient incubation of droplets in bulk. Thereby we attain a 180-fold higher number of droplets and a higher cultivation density than the microfluidic streak plate^18^, for instance (50.000 x 180 pL droplets in an array). In contrast to systems relying on agarose stabilized gel microdroplets dispersed and incubated in aqueous medium^12,42^, our method does not suffer from cross contamination of cells between the compartments or compartment and continuous phase. Although we could not use commercial FACS instruments for droplet sorting (since our droplets depend on the surrounding oil), we demonstrated a stable on-chip sorting which was implemented into a screening procedure. The screening was conducted after an extensive aerobic incubation period allowing the environmental cells to replicate and reach the physiological state for natural product synthesis before the reporter strain was injected into the compartments. Such a protocol is not feasible when double emulsions are used instead^19^.

For the cultivation of a complex bacterial community rich in recalcitrant and slow growing genera, we have demonstrated the advantages of our in-droplet cultivation strategy in comparison with standard agar plating. A more diverse subset of cultured bacteria and a two-fold higher CFU concentration compared to plates have presumably been achieved for three reasons: (i) Absence of competition in single cell confinement, (ii) the high number of incubated cells, and (iii) the new and unique cultivation conditions in our droplets during long-term dynamic droplet incubation.

It is noteworthy that the surfactant-stabilized droplet border, though forming a mechanical barrier preventing cell cross contamination, is not completely abolishing molecular transport. Small organic molecules, which are usually poorly soluble in the perfluorinated continuous phase, can leak into the continuous phase mediated by the surfactant^43^ leading to some cross talk between droplets. Although cross talk might be problematic for the detection of antibiotics, since the local concentration in the producer droplet could be diluted towards other droplets causing either false negative or false positive detection, it might present an advantage in the cultivation phase. Allowing communication between species might preserve the growth-stimulating influence of multispecies communities, which might be the key to push some of the recalcitrant species into cell propagation.

Furthermore, we were able to recover part of the increased bacterial diversity during strain isolation from droplets, exemplified by several rare genera and a candidate for a new species among relatively few isolates. Presumably, the yield of unusual isolates could be further increased by solely focusing on exceptional morphologies during colony picking, which we avoided to learn about the abundance distribution of the species diversity we obtained with the practiced isolation workflow. Here, we have isolated strains after one incubation round in droplets but alternatively, the emulsion can be broken under controlled conditions and a new droplet population started from the pooled cell suspension (data not shown). By passaging the community several times through the in-droplet cultivation, replicating organisms get enriched and the chance of strain domestication increases^16^. Furthermore, the isolation procedure enabled us to spread droplets on agar plates without mechanically disrupting cell agglomerates. Hence, also droplets that contained large cell numbers were represented by only one colony on plate. Nevertheless, the procedure exhibited a bias towards organisms abundantly found on plates due to the deposition of droplets on plates. Consequently, a combination of our approach with the microfluidic streak plate^18^ as the next step in the upscaling process could be more favorable.

We increased the practical relevance of our cultivation approach by combining it with a screening step in order to detect and specifically select droplets containing environmental cells producing compounds with antibiotic activity. By screening with two different reporter strains, we recovered multiple environmental isolates of diverse phylogenetic background that showed activity either against Gram-negative or Gram-positive bacteria with especially one *Bacillus* strain showing high antimicrobial activities. Subsequent screening approaches in combination with HR-ESI-MS/MS unveiled the antibacterial bacillaenes and antifungal gageostatins to be the causative agents. Moreover, several of those potentially antibiotic producing isolates are likely representing a new genus, which was a major goal of our study. Even though the axenic cultures could not be maintained with standard microbiological cultivation after isolation from droplets, the cells can be used for obtaining high quality genome information. Thereby our method could help to prioritize efforts during genome mining or guide future isolation campaigns.

To increase the number of rare taxa cultivated in droplets, several parameters can be altered within the presented strategy. While we used dynamic droplet incubation to continuously supply the droplet population with oxygen, the perflourinated oil carrying the dissolved gases could also be charged with nitrogen or other gases to enable a microaerophilic or even anaerobic incubation. It was recently shown that the anaerobic incubation of human microbiomes in droplets increases the species richness dramatically^21,22^. Besides encapsulating single cells, several cells could be confined per droplet creating simplified multispecies communities, which have the potential to either support growth of dependent species^44^ or to exert a change in natural product biosynthesis^45,46^.

Our cultivation and screening strategy has proven to be comprehensive and adaptable to the challenging needs of fastidious microorganisms representing interesting sources for natural products. Therefore, we envision that the bacterial diversity of numerous natural habitats, especially including environments with limited access to samples, will be explored in more depth with our strategy. The combination of bacterial strain discovery during the in-droplet cultivation phase with subsequent selection of the most auspicious candidates for natural product biosynthesis will streamline our efforts towards new scaffold structures for the development of antibiotics.

## Experimental Procedures

### Microfluidic device fabrication and operation

Microfluidic chips were designed in AutoCAD^®^ 2015 (Autodesk^®^ Corp., USA). Designs were printed by JD phototools (Great Britain) as photomasks, which were used by Biotec TU Dresden Microstructure facility to fabricate SU8 molds. PDMS (Sylgard 184, Dow Corning, Germany) replicas were obtained from molds following standard protocols^47^ and plasma bonded to microscope glass slides. For hydrophobization, Novec 1720 (3M, Germany) was introduced into the channels while heating the chips to 100 °C. At the same temperature channels designated for electrodes were filled with low melting solder (Indalloy 19, Indium Corporation of America, USA). Fluids were actuated by high precision syringe pumps (neMESYS, Cetoni GmbH, Germany) and pressure pumps (MFCS^™^-EZ, Fluigent, France). PTFE tubing (1/16″ OD, 0.25 mm ID and 0.5 mm ID, Chromophor Technologies, Germany) was used to realize fluidic connections. The device for droplet incubation was 3D-printed by i. materialise (Belgium), the complete setup was used as described before^33^ in a humid chamber. Microfluidic operations were conducted on an inverted microscope (Axio Observer Z1, Zeiss, Germany) and imaged during flow with a Pike F-032B ASG16 camera (Allied Vision). For images of stationary droplets in observation chambers a pco.edge 5.5 m (PCO AG, Germany) was employed.

### Soil sampling and extraction

Soil was collected from the nature reserve “Schwellenburg” (51° 1′ 51,9174″ latitude, 10° 57′ 14,6013″ longitude, 263 m altitude) close to Erfurt (Germany) in May 2016. The sampling site was composed of dry grass land used for leisure activities. A spot was chosen in 20 m distance to the next path. In total 12 kg of soil were collected until a depth of 10 cm after the overlaying turf was removed. The soil was transported in a sealed plastic bag and all participants wore gloves during sampling. Further processing of the soil was conducted immediately after sampling.

For the cold extracted soil extract (CESE), soil was mixed with aq. Dest. in a ratio of 1:1 (w/w). All following steps were conducted at room temperature. The mixture was stirred for 2 h at 300 rpm and then left for sedimentation for 2 h. Further insoluble particles in the decanted supernatant were removed by centrifugation at 15970 x g for 20 min. The resulting supernatant was filtered twice through cotton wool, aided by applying vacuum. For final sterilization the filtrate was filtered through bottle-top-filters with sterile filter membranes (pore size 0.2 μm) and subsequently aliquoted. Aliquots were stored at −20 °C.

To extract the bacterial soil community, 20 g of the freshly sampled soil spread onto a flat glass tray were dried for 24 h at 37 °C. The dried soil was grinded in a mortar, while stones and macroscopic plant material were removed. In 90 mL CESE, 10 g of the milled soil and 25 glass beads (diameter 2-6 mm) were suspended and shaken at 200 rpm for 2 h at 28 °C. Subsequently the suspension was sonicated twice at 50 W for 1 min. One half of the suspension was filtered through a cell strainer (40 μm pore size) aided by applying vacuum. The other half was left for particle sedimentation at room temperature. After 1 h the supernatant was decanted. Aliquots of both cell suspensions were stored at −20 °C. Upon usage the suspension rich in soil particles was mixed with the suspension containing fewer particles in a ratio of 2:35 (v/v).

### Microorganisms and culture conditions

The soil community was cultivated in medium containing 50 % (v/v) CESE, 6 % (v/v) supernatant of soymannit medium (20 g/L soy coarse meal [Schkade Landhandel GmbH, Germany], 20 g/L mannitol [Merck KGaA, Germany], deionized water, pH adjusted to 6.5, sterilization 35 min at 121°C) and 44 % (v/v) deionized water. For agar plates 2 % (w/v) agar was autoclaved with appropriate amounts of water and soy-mannitol supernatant at 121 °C for 20 min. After cooling to 56 °C, CESE was added and plates were poured. For cultivation on plates the soil community was diluted with 0.9 % (w/v) NaCl solution in a serial dilution until 10^-4^; 50 μl of the respective dilution were spread on agar plates with Drigalski spatulas. For droplet generation the soil community was suspended in medium without dilution. The positive control comprised a spore mixture of known *Streptomyces* strains (see SI) in equal concentrations (8 x 10^5^ spores/mL) in the medium described above. All samples containing the soil community and respective controls were incubated at 20 °C.

For the inhibition assays in droplets the reporter strains *Escherichia coli* (*E. coli* JW1982^48^ with plasmid pMPAG6 - P_T5/lacO_:mCherry) and *Bacillus subtilis* (*B. subtilis* 168 *amyE::hy-mKATE:CM*^49^) were used. They were cultivated in terrific broth containing 1.2 % (w/v) tryptone (Bacto Tryptone, BD Bioscience, Belgium), 2.4% (w/v) yeast extract (Bacto Yeast Extract, BD Bioscience, Belgium), 0.4% (v/v) glycerol (Roth, Germany), 0.17 M KH_2_PO_4_ (Merck, Germany) and 0.72 M K_2_HPO_4_ (Merck, Germany) in tap water (pH adjusted to 7.2 with NaOH) + 1% (w/v) glucose (VWR International, USA) containing 100 μg/mL ampicillin (*E. coli*) or 5 μg/mL chloramphenicol (*B. subtilis*) at 37 °C, 200 rpm/min for 16 h. New cultures were inoculated to a start OD_600_ of 0.1 and cultivated without selection markers until midexponential phase. Before pico-injection of reporter cells into droplets, cells were pelleted and resuspended in 2.5x terrific broth + 1% (w/v) glucose to a final OD_600_ of 4. In case of *E. coli* 0.5 mM IPTG were added.

### Comparison of cultivation in droplets and on agar plates

Microfluidic droplets were generated at a flow focusing unit, using Novec HFE7500 (3M, Germany) with 0.5% Pico-Surf 1 (Dolomite, UK) as continuous phase. Droplets were generated usually at a frequency of 1500 Hz and with a volume of 200 pL. One droplet population, which was incubated in one droplet incubator, comprised approximately 9 x 10^6^ droplets. Droplet populations were generated with 8 replicates and compared to 8 plate sets. One plate set consisted of 8 plates with 10^-2^ dilution of the soil community and 7 plates with 10^-3^ dilution. For each cultivation technique three positive controls were prepared and incubated along with the soil community samples. After the incubation time of 28 days, microcultures of one droplet population were pooled by fusing the emulsion with 1H, 1H, 2H, 2H-Perfluoro-1-octanol (Sigma, USA). The upper aqueous phase including the cells was transferred into a new vessel and the cells were pelleted. From plates, colonies were pooled by suspending the cells in 2 mL 0.9% NaCl per plate with a cell scraper (Corning^®^, VWR, Germany).

For DNA extraction 150 mg of sterilized glass beads (0.1-0.5 mm diameter, in case) were added along with one volume of lysis buffer (100 mM Tris-HCl (pH 8), 20 mM Na-EDTA, 1.5 M NaCl, 1.2% (w/v) Triton X-100) to the cell pellets. In case of cell pellets derived from plates, the double amount of glass beads and 3 ceramic beads (0.5 cm diameter) were used. Cells were lysed in a FastPrep-24 (MP Biomedicals, Germany) two times for two minutes at a speed of 6 m/s. Afterwards, 0.3 volumes of 20% (w/v) SDS (Roth, Germany) were added and precipitates removed by centrifugation. The cell lysate was extracted with one volume of phenol:chloroform:isoamylalcohol (25:24:1, Roth, Germany). To remove phenol traces, the samples were further extracted three times with one volume of chloroform:isoamylalcohol (24:1, Roth, Germany). Finally, the DNA was precipitated with ethanol and dissolved in NE Buffer. To remove humic acids, the DNA was further purified with custom made Q Sepharose columns as described by Mettel et al.^50^. See supplemental information for more details on precipitation and purification via Q Sepharose.

The V3-V4 region of the 16S rRNA gene was amplified using the primers 314F and 758R^51^ with Illumina adapters at the 5’ end. The Q5 polymerase (New England Biolabs, Germany) was used in a 50 μl reaction after pretreatment with DNase I (New England Biolabs, Germany) according to the manufacturer’s protocol (see SI for more details). The settings for the PCR were pre-denaturation at 98 °C for 3 min, 30 cycles of denaturation at 98 °C for 30 s, annealing at 55 °C for 40 s, elongation at 72 °C for 1 min and final elongation at 72 °C for 5 min. DNA libraries were constructed from the PCR products following the standard Illumina protocol. Amplicons were sequenced on an Illumina MiSeq system with 200 bp paired end by IIT Biotech (Germany).

### Cell isolation from microfluidic droplets

Droplets were generated and incubated with the soil community as described above. After one month, droplets were re-injected from the incubator into a microfluidic chip, where they were spaced to a frequency of 10-50 Hz. All droplets were directed to a glass capillary (TSH, 100 μm ID, 360 μm OD, Molex, USA) which led to a positioning system (neMAXYS 200, cetoni GmbH, Germany). The tip of the capillary was attached to a z-steerable arm of the system, while an agar plate was positioned below on a x-y-steerable platform. Thereby droplets were continuously deposited in a spiral pattern on an agar plate over which the outlet of the capillary was moved at 1 mm/s. Plates were incubated for 15 days at 20 °C, after which colonies were picked and streaked onto new agar plates. By re-streaking multiple times pure isolates were obtained, of which DNA was extracted using the QIAamp DNA Mini Kit (Qiagen, Germany). For isolate characterization the whole 16S rRNA gene was amplified in 50 μl reaction using the primers 27F (AGA GTT TGA TCM TGG CTC AG) and 1492R (CGG TTA CCT TGT TAC GAC TT) and the PrimeSTAR GXL polymerase (Takara Bio, USA). Before amplification the polymerase was treated with DNase I. The PCR was carried out as follows: predenaturation at 98 °C for 30 s, 35 cycles of denaturation at 98 °C for 20 s, annealing at 55 °C for 40 s, elongation at 68 °C for 30 s and final elongation at 68 °C for 3 min. The Sanger sequencing of forward and reverse strand was done by Macrogen (Netherlands).

### Screening for antimicrobial compounds in droplets

Droplets inoculated with the soil community and incubated for 28 days were re-injected into a microfluidic chip at a frequency of 200 Hz. The chip contained a pico-injection structure similar to structures described by Abate et al.^52^. By applying an alternating electrical field through a function generator (AFG-2005, GW Instek, China) and a high-voltage amplifier (model 2210-CE, Trek, USA) with a final amplitude of 20 V, a frequency of 20 kHz and a duty cycle of 50%, a cell suspension containing the reporter strain was pico-injected into each droplet. Droplets were subsequently guided into a new incubator and soil organisms and reporter cells were co-incubated for 20 h at 28 °C inside droplets by applying dynamic droplet incubation. Finally droplets were re-injected into a droplet sorting structure^53^, where the fluorescence protein of the reporter strains was excited with a Laser (488 nm diode laser, Lasos, Germany) and detected with a photomultiplier module (H10721-20, Hamamatsu Photonics UK Limited, UK). Droplets exceeding a pre-defined intensity threshold were sorted via dielectrophoresis by a burst signal of 430 mV (1000x amplified), 30 cycles, 6 kHz, 50% duty cycle. Software was custom-made and written in LabVIEW (v2015). Sorted droplets were discarded while remaining droplets were collected and distributed on agar plates. Plates were incubated, and colonies were picked, purified and characterized as described above.

### Data analysis of sequence data

Forward and reverse reads of the amplicon data were merged by FLASH^54^. Primers were removed with cutadapt^55^ allowing 20 % error. For quality trimming, sickle^56^ was used with a minimum quality threshold of 20. Reads were further processed following the MiSeq SOP (https://www.mothur.org/wiki/MiSeq_SOP) in Mothur^57^. To detect chimera the implemented uchime algorithm was used in combination with the SILVA reference database (release 128)^58^, which was also used to taxonomically classify the reads. For clustering into operational taxonomic units (OTU) the vsearch method “abundance based greedy clustering” and a distance cutoff of 0.03 were employed. Read numbers across all samples were subsampled to 88435 sequences, to obtain a better comparability. The OTU table was further processed and visualized in R and with the R package phyloseq^59^. OTUs were excluded, when they were not represented more than three times in at least 10% of the samples. Indices for alpha diversity are shown in Supplemental Information for subsampled and not subsampled data.

Consensus 16S rRNA gene sequences of axenic cultures were assembled from forward and reverse reads with SeqTrace^60^ (v0.9.0) applying a Needleman-Wunsch alignment algorithm and a quality cut off for base calls of 30. After automatic trimming until 20 out of 20 bases were correctly called, consensus sequences were examined and curated manually. Consensus sequences of the nearly full-length 16S rRNA gene were aligned with SILVA Incremental Aligner (SINA)^61^ (v1.2.11). Phylogenetic relations were deduced by reconstructing phylogenetic trees with ARB^62^ (v6.0.6) using the ‘All species living tree project’ database^63^ (release LTPs128, February 2017). Sequences were added into the LTP type strain reference tree using ARB parsimony (Quick add marked) and alignment was corrected manually. Phylogenetic tree calculation with all family members was based on maximum-likelihood algorithm using RAxML^64^ (v7.04) with GTR-GAMMA and rapid bootstrap analysis, maximum-parsimony method using DNAPARS^65^ (v3.6), and neighbour-joining with Jukes–Cantor correction.

### Secondary metabolite production

For the identification of natural products, *Bacillus* sp. strain D121-0906-b3-2-1 was inoculated in diverse media (MGY+M9^66^ and soy mannit medium + CESE) at OD_600_ 0.1 and grown at 160 rpm, 28 °C for 3-8 days. Extraction took place with 1:1 volume of ethyl acetate overnight or with addition of XAD-2 to the culture broth for 30 minutes followed by elution twice with 100% methanol for 30 minutes. The organic phase was dried with anhydrous sodium sulfate and concentrated under reduced pressure. Residues were dissolved in a small volume of methanol and measured at LC/MS. Fragmentation patterns were monitored *via* tandem mass spectrometry (MS/MS).

#### LC/MS

Exactive Orbitrap High Performance Benchtop LC-MS (Thermo Fisher Scientific) with an electron spray ion source and an Accela HPLC System, C18 column (Betasil C18, 150 x 2.1 mm, Thermo Fisher Scientific), solvents: acetonitrile and distilled water (both supplemented with 0.1% formic acid), flow rate: 0.2 mL/min; program: hold 1 min at 5% acetonitrile, 1–16 min 5–99% acetonitrile, hold 15 min 99% acetonitrile, 19–20 min 99% to 5% acetonitrile, hold 11 min at 5% acetonitrile.

#### MS/MS

QExactive Orbitrap High Performance Benchtop LC-MS (Thermo Fisher Scientific) with an electron spray ion source and an Accela HPLC System, C18 column (Accucore C18 2.6 μm, 100×2.1 mm, Thermo Fisher Scientific), solvents: acetonitrile and distilled water (both supplemented with 0.1% formic acid), flow rate: 0.2 mL/min; program: hold 1 min at 5% acetonitrile, 1–10 min 5–98% acetonitrile, hold 12 min 98% acetonitrile, 22–22.1 min 98% to 5% acetonitrile, hold 7 min at 5% acetonitrile.

The metabolic profiles of the acquired fractions were monitored with HR-ESI-LC/MS.
Bacillaene A: measured 579.3454 [M−H]^-^, calculated 579.3434 C_34_H_47_O_6_N_2_
Bacillaene B: measured 741.3990 [M−H]^-^, calculated 741.3962 C_40_H_57_O_11_N_2_
Gageostatin A: measured 1040.6859 [M+H]^+^, calculated 1040.6859 C_52_H_94_O_14_N_7_
Gageostatin B: measured 1054.7014 [M+H]^+^, calculated 1054.7015 C_53_H_96_O_14_N_7_
Gageostatin C: measured 1008.6594 [M+H]^+^, calculated 1008.6597 C_51_H_90_O_13_N_7_

## Acknowledgements

The authors thank Karin Burmeister for excellent technical assistance and Dr. Àkos Kovács for kindly providing the *B. subtilis* strain. This work has been supported by the Thuringian Ministry of Education, Science and Culture and the European Fond for Structural Development (project MicroInfra no. 13008-715, CCI-Code 2007DE161PO001), the Thuringian Ministry of Economy, Labor and Technology (project DropCode, no. 2014FE9037), the German Center for Infection Research funded by the Federal Ministry of Education and Research (project no. TTU 09.811), and the Jena School for Microbial Communication (JSMC) funded by the German Excellence Initiative.

