## Supplemental Information for "Highly parallelized microfluidic droplet cultivation and prioritization on antibiotic producers from complex natural microbial communities"

### List of Videos

1 Picoinjection of reporter cells to droplets inoculated and incubated with the soil community. . . . . 12
2 Sorting of droplets based on their fluorescence intensity in the red channel. . . . . 12

### List of Tables

1 Positive control for droplet-plate comparison . . . . . 2
2 Screenings for antibiotics in droplets with 2 different reporter strains and 2 different media. . . . . 12

### Material and Methods

**Table S 1** – Positive control for droplet-plate comparison

| Species | ID |
|---|---|
| *Streptomyces griseus* | ST036300 |
| *Streptomyces hygroscopicus* | HKI0016 |
| *Streptomyces noursei* | JA03890 |
| *Streptomyces netropsis* | IMET43883 |
| *Streptomyces collinus* | IMET43780 |

#### Ethanol precipitation of extracted metagenomic DNA

DNA samples were mixed with 0.1 volume 3 M sodium acetate, 0.001 volume of 20 mg/mL glycogen and 2.5 volumes of 98% (v/v) cold ethanol. Samples were stored at -20 °C for at least 16 h before they were centrifuged for 1 h at 4 °C and 25000 x g. The supernatant was removed and the remaining pellet washed with 1 mL 70% (v/v) ethanol. The samples were centrifuged and decanted again. Traces of ethanol were removed by drying the pellet for 5 min under vacuum. Afterwards the DNA was rehydrated in NE Buffer (5 mM Tris/HCl, pH 8.5) and stored at 4 °C.

#### Further purification of metagenomic DNA via Q Sepharose columns

For each DNA sample 600 µL of Q Sepharose (Q Sepharose High Performance, GE Healthcare, USA) were aliquoted in 1.5 mL reaction vessels and washed 4 times in 1 mM potassium phosphate buffer (136 mg/L $KH_2PO_4$, 228 mg/L $K_2HPO_4$ x 3 $H_2O$). After washing the Q Sepharose pellet was resuspended in 300 µL of the same buffer and transferred into a centrifugal filter unit (Durapore-PVDF 0.45 µm, Merck Millipore Ltd, Germany). The self-made column was packed by centrifugation, buffer that passed through the filter membrane was discarded. In the next step the DNA sample dissolved in NE Buffer was pipetted onto the column and centrifuged for 10 s at 3000 x g, thereby DNA and humic acids were binding to the Q Sepharose. The DNA was eluted by adding 80 µL of a 1.5 M NaCl solution which was also pushed through the column by centrifugation at 3000 x g for 10 s. The elution step was repeated until the clearly visible brown band of humic acids reached the filter membrane, which usually happened after 4 repetitions. The DNA in the eluate was desalted by binding it to a silica membrane using the DNA Clean & Concentrator Kit (Zymo Research, Germany) following the manufacturer's protocol.

### DNase I pretreatment of polymerases

The polymerases were pretreated with DNase I in order to remove bacterial DNA traces that originated from the enzyme production processes. The necessary amount of polymerase stock was diluted in a ratio of 1:6 in PCR grade water. 10x DNase I buffer and DNase I were added according to the manufacturer's protocol. The mixture was incubated 10 min at 37 °C and inactivated 10 min at 75 °C. After cooling down, the mixture was added to the master mix, which was subsequently aliquoted for the following PCR.

### Image analysis for fluorescence images of stationary droplets in observation chambers

Images were analyzed as virtual stack in fiji[1]. The copy of a bright-field image was segmented in order to locate droplet borders and define regions of interest (ROIs) within the droplet borders. With a custom-made macro the contrast of the image was enhanced until 1% of the pixels reached saturation. To reduce noise a median filter was applied before the local threshold method phansalker was used with a radius of 15 pixels to binarize the image. Remaining black foreground structures, which mainly represented the droplet borders, were dilated in lateral direction by 5 pixels. To recognize the inside of the droplet borders as ROIs the binary image was inverted, resulting in droplet borders now becoming white background, thereby allowing to run the function "analyze particles". The radius of each detected ROI was reduced by 5 pixels to avoid overlap of the ROI with droplet borders. Not recognized droplets were manually added to the list of ROIs. The finalized list of ROIs was used to read out the average grey values in original images of bright-field, dark-field and fluorescence channels. Thereby tables were created with intensities for all fluorescence channels for every analyzed droplet. The tables were further processed and visualized in R.

### Additional Results

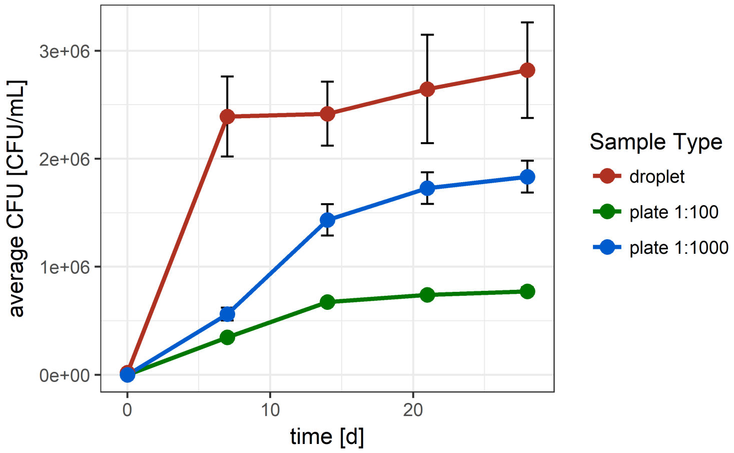

**Figure S 1** – Average CFU concentration over time at the different cultivation methods. Displayed is the average CFU concentration over time obtained for the extracted soil community by cultivating the soil community in droplets or on plates using the same media composition and equal incubation time. Error bars are showing the standard error of the mean.

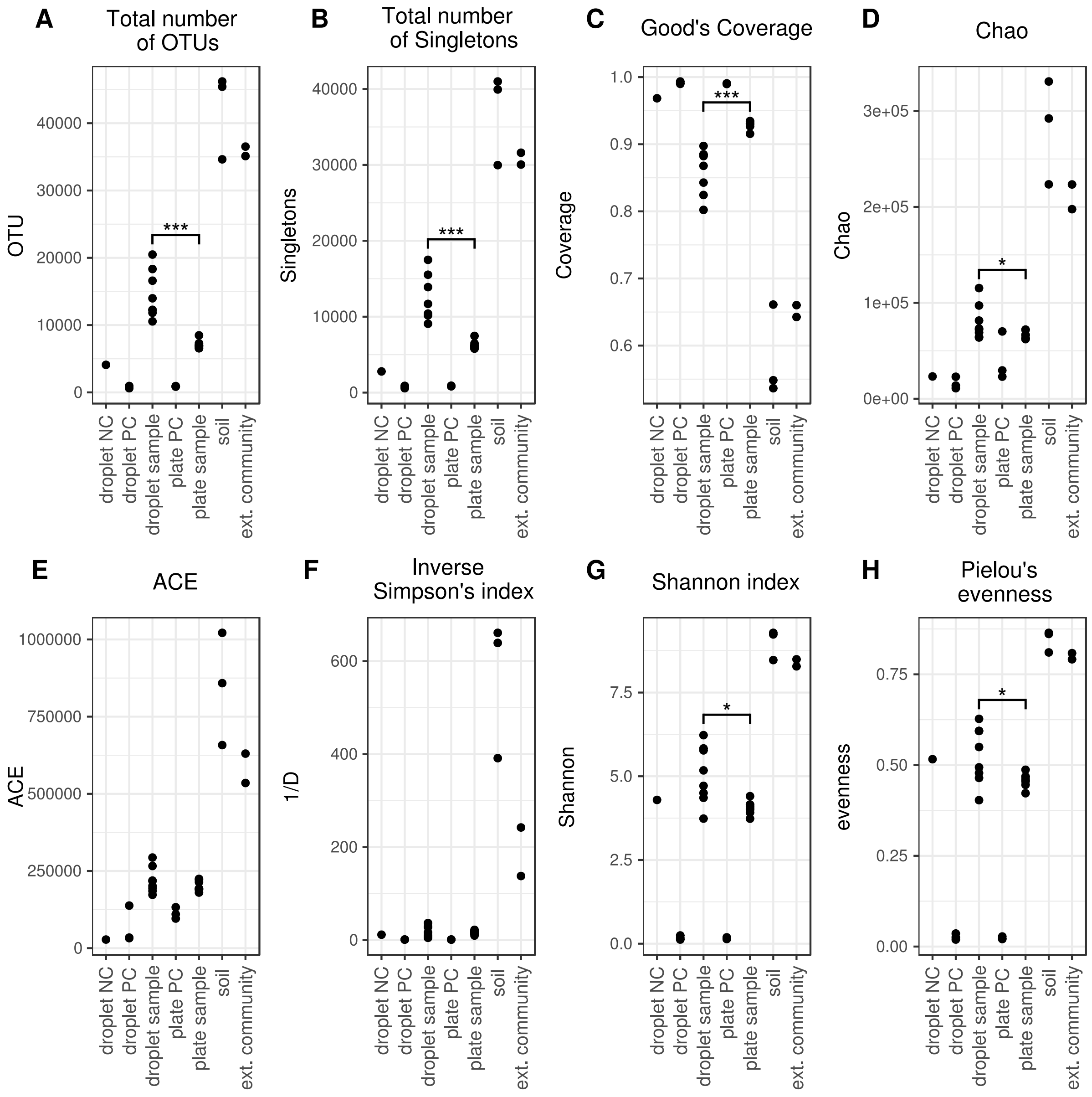

**Figure S 2** – Alpha diversity indices for the read abundance normalized 16S rRNA datasets. Means between droplet sample and plate sample were tested by Wilcoxon Rank-Sum test. Cut-off for significance was $p < 0.05$. *** significant with $p < 0.001$, * significant with $p < 0.05$.

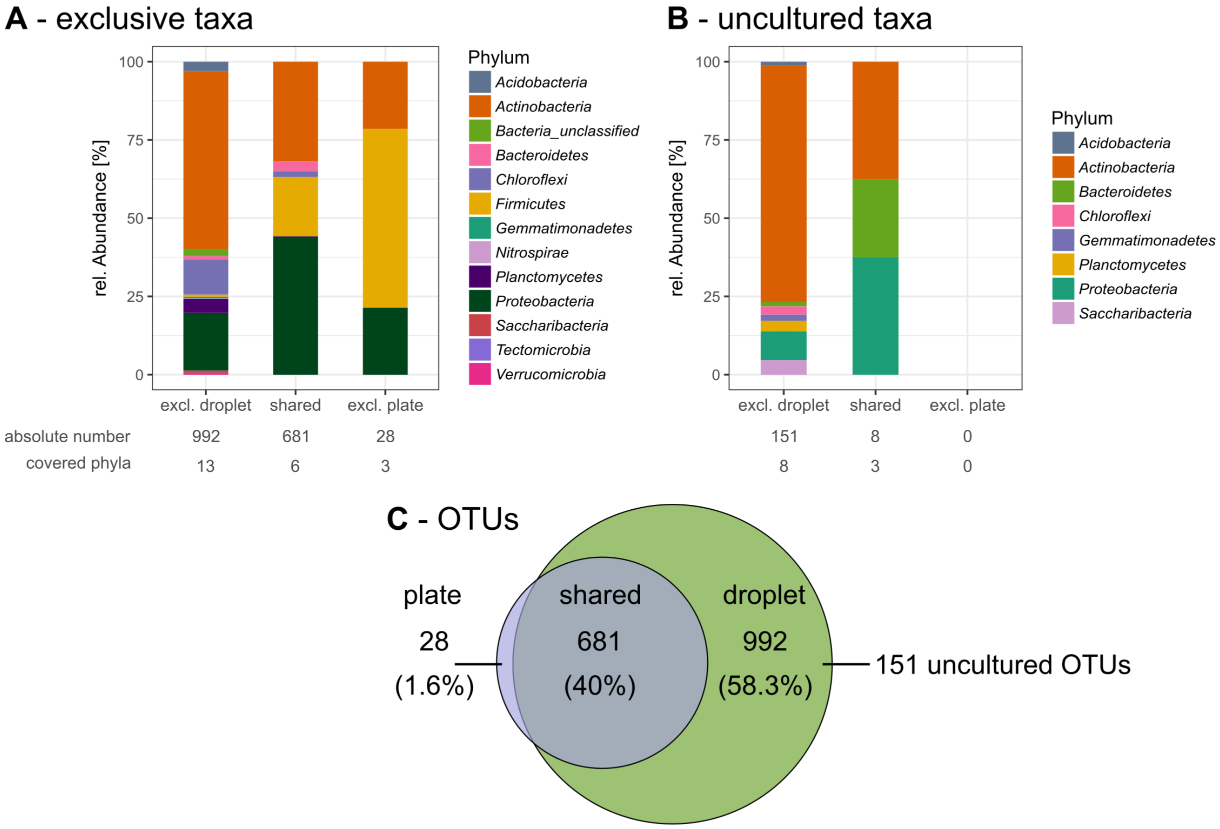

**Figure S 3** – A) Overview of taxa that were exclusively found in droplet samples and/or plate samples ith taxonomic classification on the phylum level. B) Overview of the exclusive taxa that were classified as uncultured found in droplet and/or plate samples. Taxa that were unclassified on the genus level and likely belong into the uncultured category were not considered for this comparison. C) Explanation of the terminology. Exclusive taxa were either found only in droplet samples or plate samples. Shared taxa were found in both sample types.

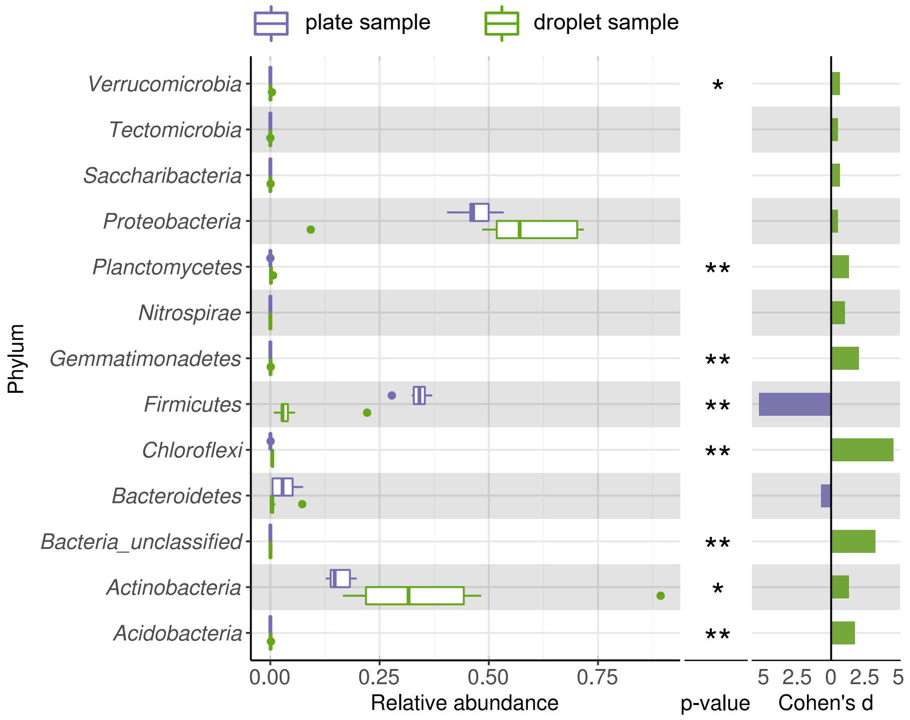

**Figure S 4** – Comparing taxonomic classification on phylum rank for replicating cells in agar plate and in-droplet cultivation. Boxplots depict the distribution of relative abundances for phyla in droplet and plate cultivation samples. Relative abundances of OTUs were agglomerated on the phylum level. Displayed are 13 assigned phyla covering 100% of all sequences. Means of relative abundances were compared by Wilcoxon Rank-Sum test for each genus ($\alpha$=0.05), applying Holm-Bonferroni correction for multiple comparisons. ** significant with $p < 0.01$, * significant with $p < 0.05$. As effect size Cohen's d was computed and plotted as bars, to indicate which differences are practically relevant. The direction and the color of the bars depend on the sample type in which the larger mean was found (blue – larger mean in plate samples, green – larger mean in droplet samples).

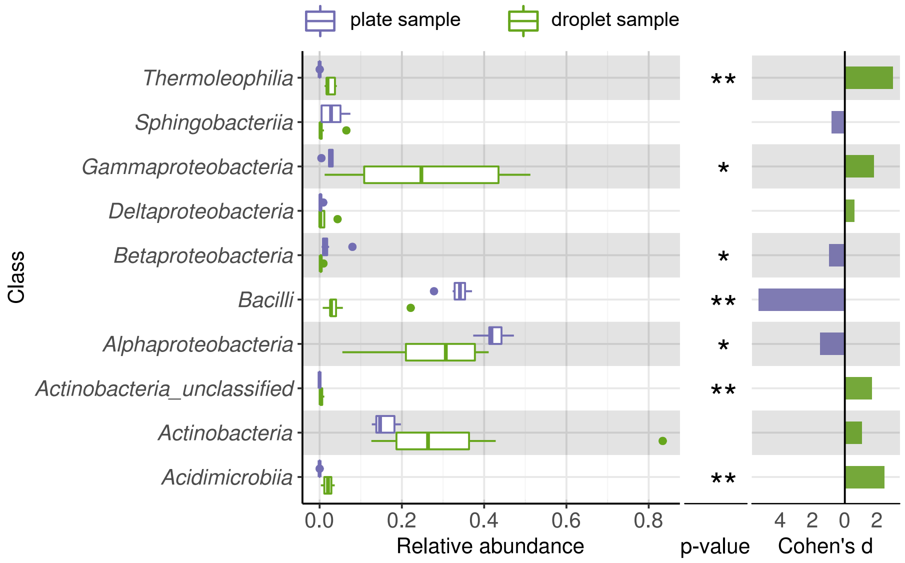

**Figure S 5** – Comparing taxonomic classification on class rank for replicating cells in agar plate and in-droplet cultivation. Boxplots depict the distribution of relative abundances for classes in droplet and plate cultivation samples. Relative abundances of OTUs were agglomerated on the class level. Displayed are the 10 most abundant classes of in total 41 assigned classes covering 99.42% of all sequences. Means of relative abundances were compared by Wilcoxon Rank-Sum test for each genus ($\alpha$=0.05), applying Holm-Bonferroni correction for multiple comparisons. ** significant with $p < 0.01$, * significant with $p < 0.05$. As effect size Cohen's d was computed and plotted as bars, to indicate which differences are practically relevant. The direction and the color of the bars depend on the sample type in which the larger mean was found (blue – larger mean in plate samples, green – larger mean in droplet samples).

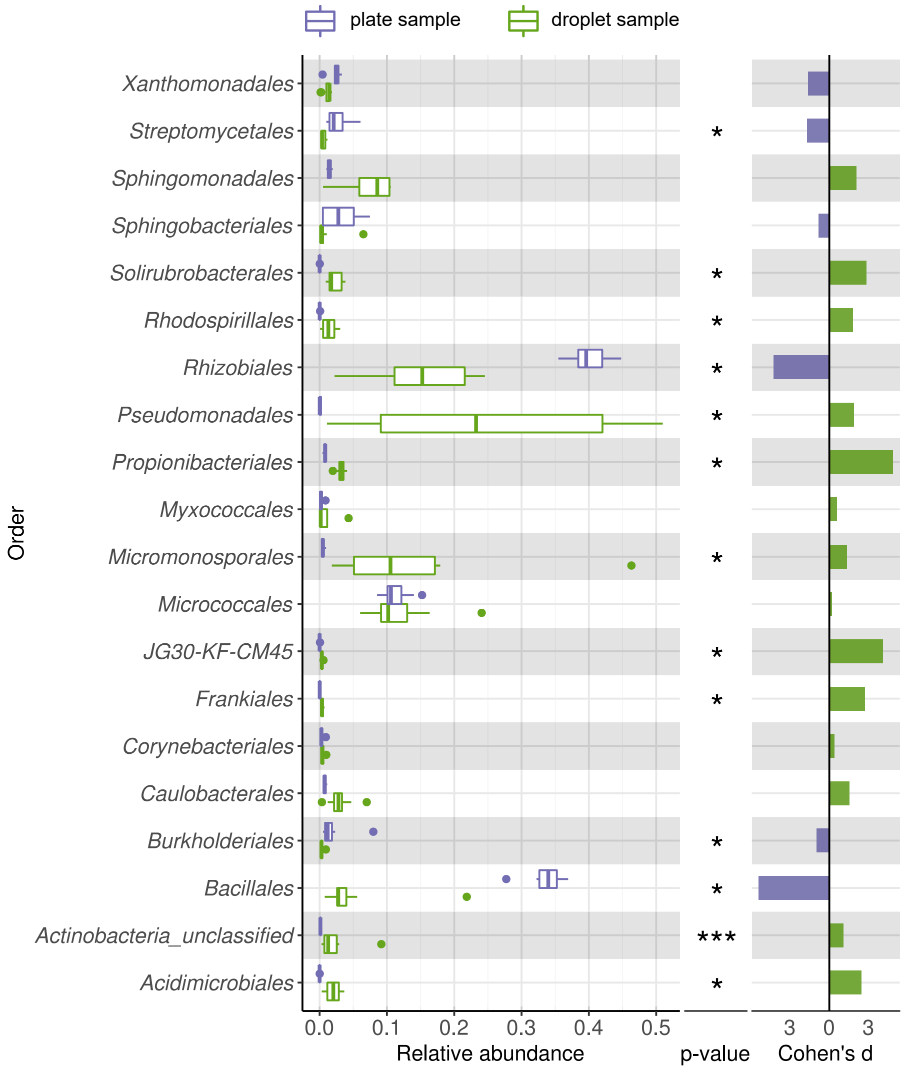

**Figure S 6** – Comparing taxonomic classification on order rank for replicating cells in agar plate and in-droplet cultivation. Boxplots depict the distribution of relative abundances for orders in droplet and plate cultivation samples. Relative abundances of OTUs were agglomerated on the order level. Displayed are the 20 most abundant orders of in total 72 assigned orders covering 99.03% of all sequences. Means of relative abundances were compared by Wilcoxon Rank-Sum test for each genus ($\alpha$=0.05), applying Holm-Bonferroni correction for multiple comparisons. *** significant with $p < 0.001$, ** significant with $p < 0.01$, * significant with $p < 0.05$. As effect size Cohen's d was computed and plotted as bars, to indicate which differences are practically relevant. The direction and the color of the bars depend on the sample type in which the larger mean was found (blue – larger mean in plate samples, green – larger mean in droplet samples).

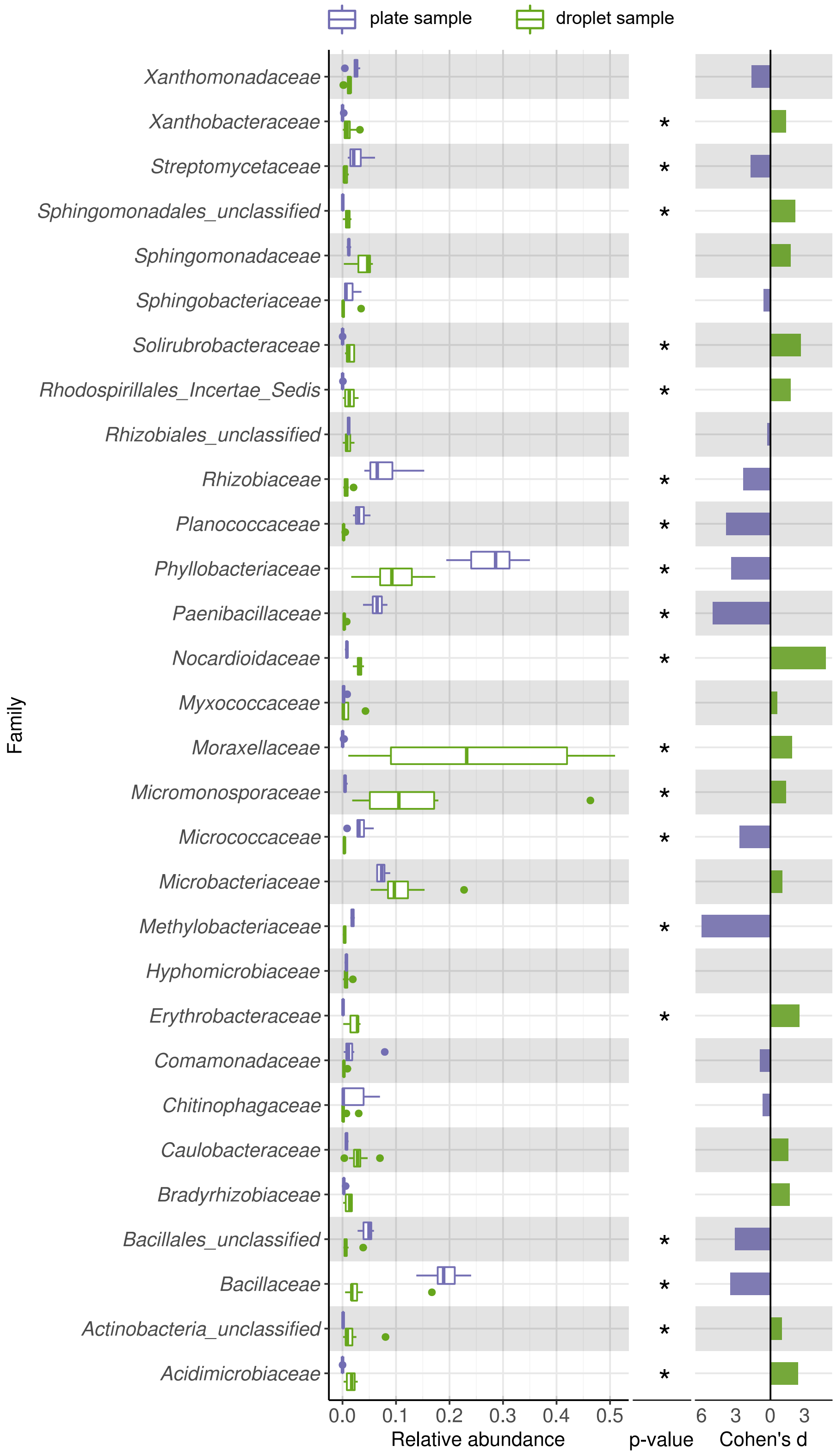

**Figure S 7** – Comparing taxonomic classification on family rank for replicating cells in agar plate and in-droplet cultivation. Boxplots depict the distribution of relative abundances for families in droplet and plate cultivation samples. Relative abundances of OTUs were agglomerated on the family level. Displayed are the 30 most abundant classes of in total 144 assigned families covering 96.37% of all sequences. Means of relative abundances were compared by Wilcoxon Rank-Sum test for each genus ($\alpha$=0.05), applying Holm-Bonferroni correction for multiple comparisons. ** significant with $p < 0.01$, * significant with $p < 0.05$. As effect size Cohen's d was computed and plotted as bars, to indicate which differences are practically relevant. The direction and the color of the bars depend on the sample type in which the larger mean was found (blue – larger mean in plate samples, green – larger mean in droplet samples).

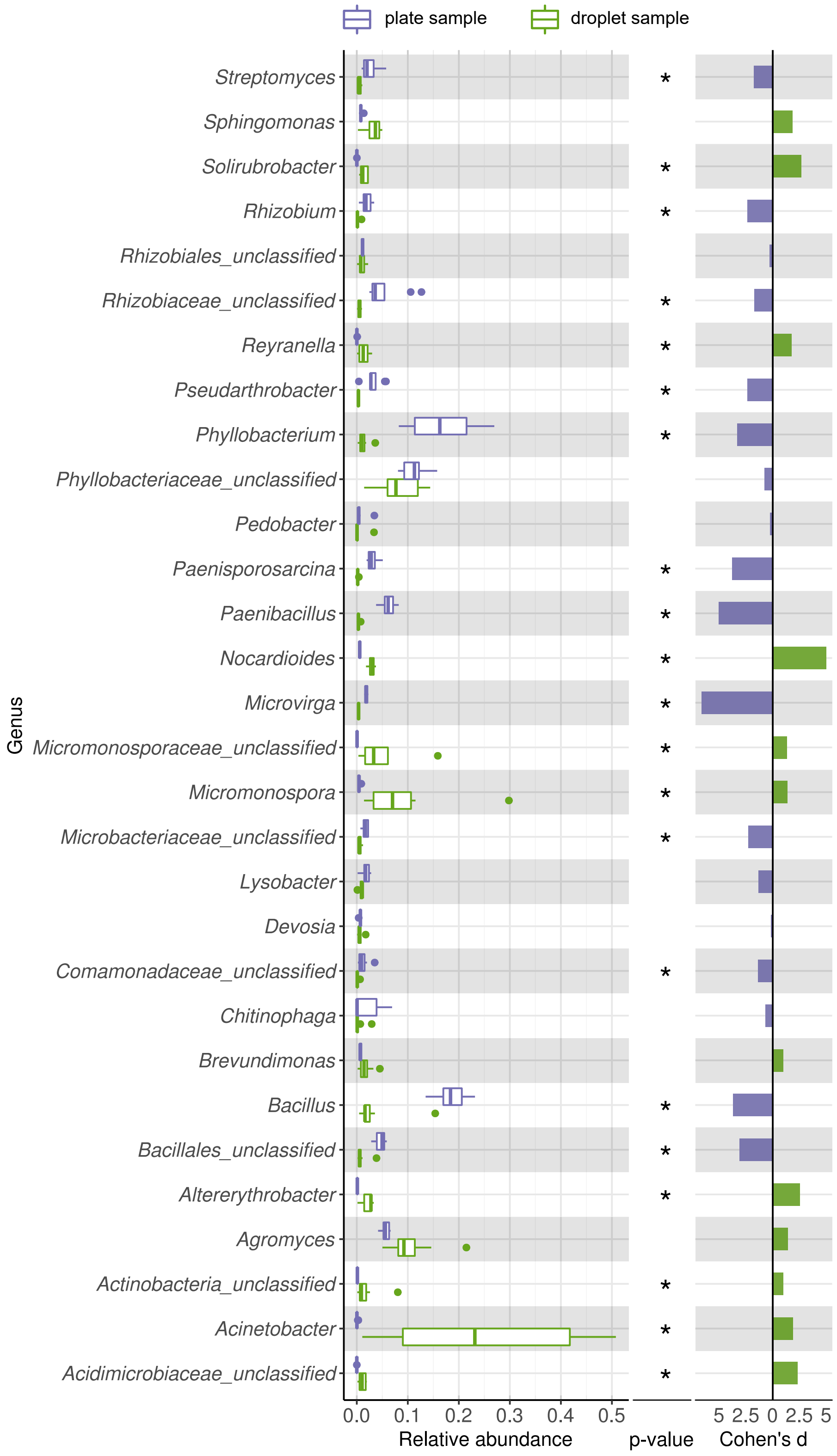

**Figure S 8** – Comparing taxonomic classification on genus rank for replicating cells in agar plate and in-droplet cultivation. Boxplots depict the distribution of relative abundances for genera in droplet and plate cultivation samples. Relative abundances of OTUs were agglomerated on the genus level. Displayed are the 30 most abundant genera of in total 240 assigned genera covering 89.54% of all sequences. Means of relative abundances were compared by Wilcoxon Rank-Sum test for each genus ($\alpha$=0.05), applying Holm-Bonferroni correction for multiple comparisons. ** significant with $p < 0.01$, * significant with $p < 0.05$. As effect size Cohen's d was computed and plotted as bars, to indicate which differences are practically relevant. The direction and the color of the bars depend on the sample type in which the larger mean was found (blue – larger mean in plate samples, green – larger mean in droplet samples).

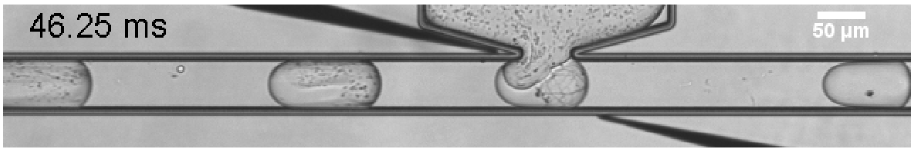

**Video S 1** – Picoinjection of reporter cells to droplets inoculated and incubated with the soil community. Images were taken with 800 frames/s at 5x magnification in bright-field illumination. Images are played with 7 frames/s. In the field of view is the channel, which is guiding the reinjected droplets through the electrical field applied between the black electrodes (only tips are visible). The surfactant stabilized water oil interphase is destabilized by the electrical field resulting in fusion of the aqueous phase coming from the top channel and containing reporter cells and the aqueous droplet. When droplets leave the electrical field the interphase is stable again.

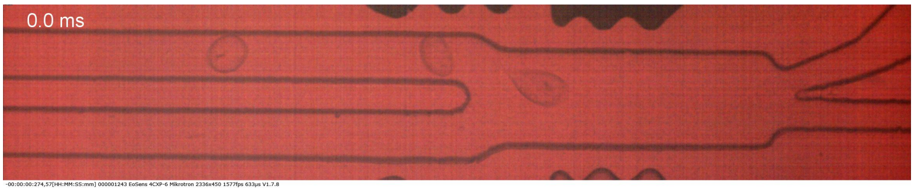

**Video S 2** – Sorting of droplets based on their fluorescence intensity in the red channel. Images were taken with 1577 frames/s at 5x magnification. Every third image is played with 10 frames/s, resulting in 52-fold reduction in playback speed. In the field of view is the droplet sorting structure. Droplets with high red fluorescence intensity are pulled with an electrical field into the upper channel. Droplets with low red fluorescence intensity leave the chip through the lower outlet.

**Table S 2** – Screenings for antibiotics in droplets with 2 different reporter strains and 2 different media.

| | **Medium** | **Reporter strain** | **Nb. of isolates** | **Nb. of isolates characterized** |
|---|---|---|---|---|
| 1 | 50% CESE, 6% soy mannit medium | *E. coli* | 67 | 17 |
| 2 | 50% CESE, 6% malt medium | *E. coli* | 132 | 68 |
| 3 | 50% CESE, 6% soy mannit medium | *B. subtilis* | 180 | 63 |
| 4 | 50% CESE, 6% malt medium | *B. subtilis* | 78 | 9 |

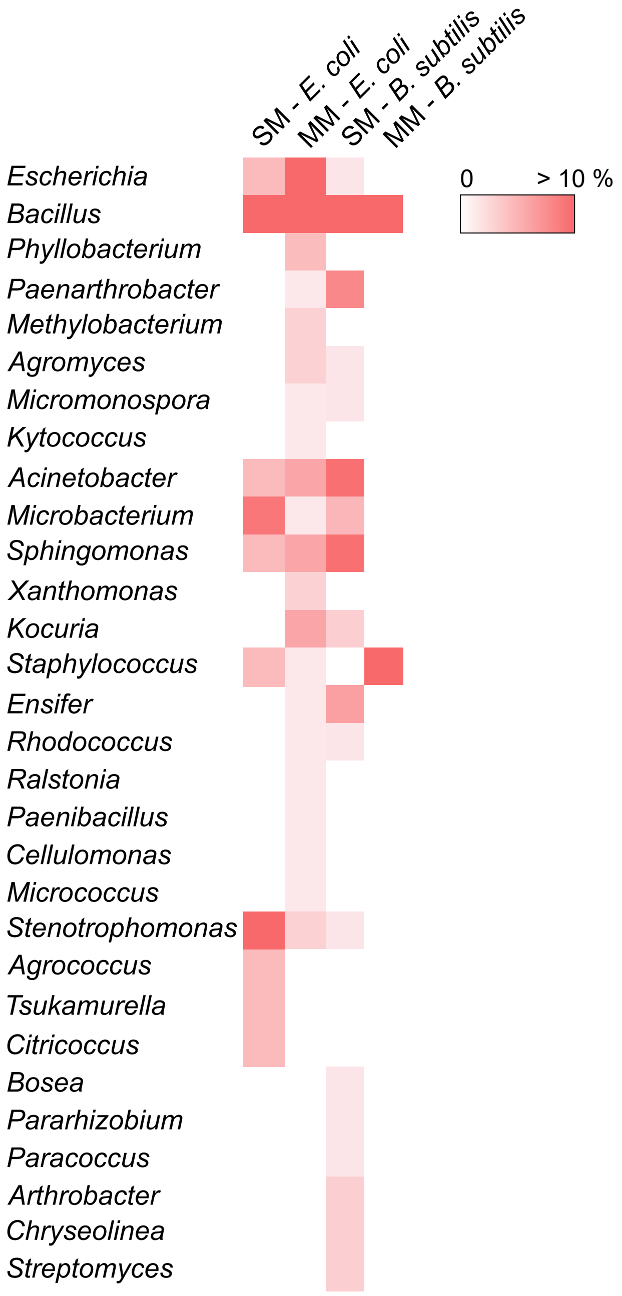

**Figure S 9** – Abundance of genera isolated in four different screening experiments for antimicrobial activity depicted as heatmap. All identified genera are listed with their observed relative abundance in the corresponding experiment. White squares were assigned when the genus was not represented by isolates of the experiment. SM denotes soy mannit medium, MM denotes malt medium. Absolute numbers of characterized isolates were: SM - *E. coli*: 17, MM - *E. coli*: 68, SM - *B. subtilis*: 63, MM - *B. subtilis*: 9.

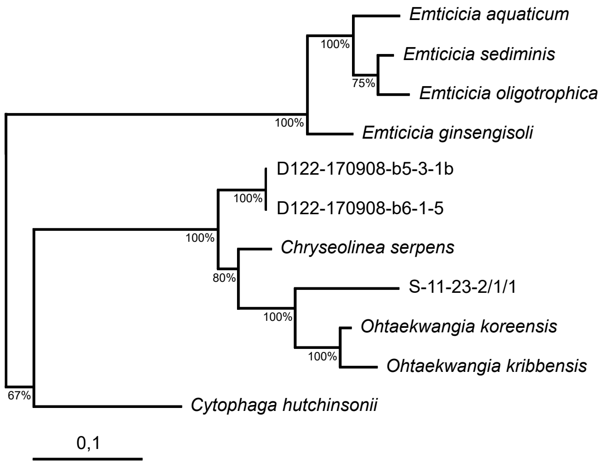

**Figure S 10** – Maximum likelihood tree based on nearly full length 16S rRNA gene sequence. The tree shows the phylogenetic placement of isolates among type strains, belonging to the Cytophagaceae. Bootstrap values (100 resamplings) are indicated at nodes. Isolate names starting with D were obtained in targeted isolation, while S-11-23-2/1/1 was isolated in the untargeted approach.

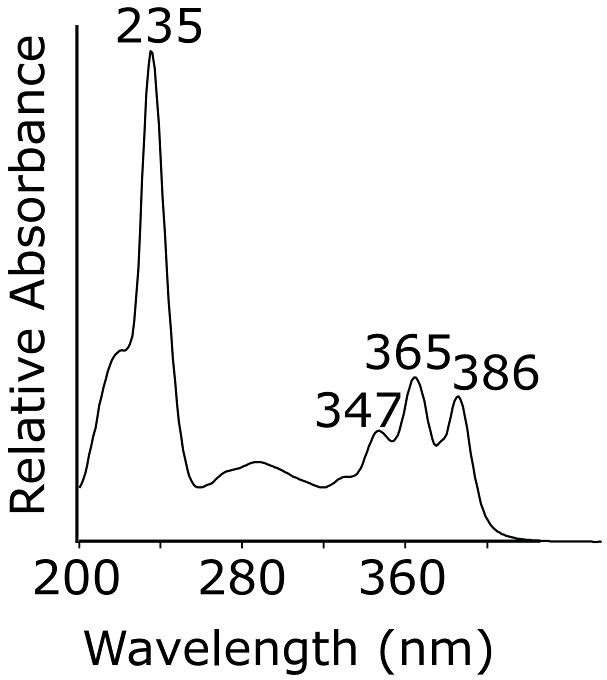

**Figure S 11** – Examplary UV absorption spectrum for bacillaenes.

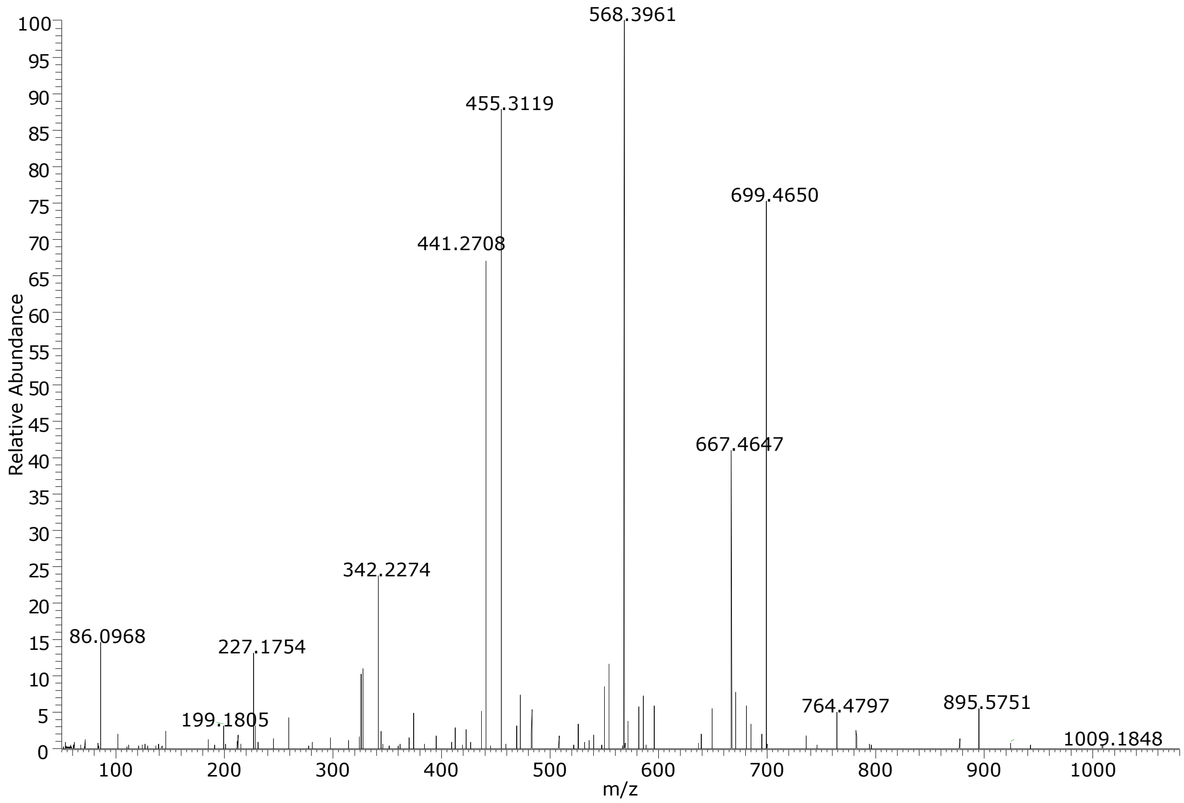

**Figure S 12** – MS/MS fragmentation pattern of $[M+H]^+$ 1040.6859 (gageostatin A) in positive mode.

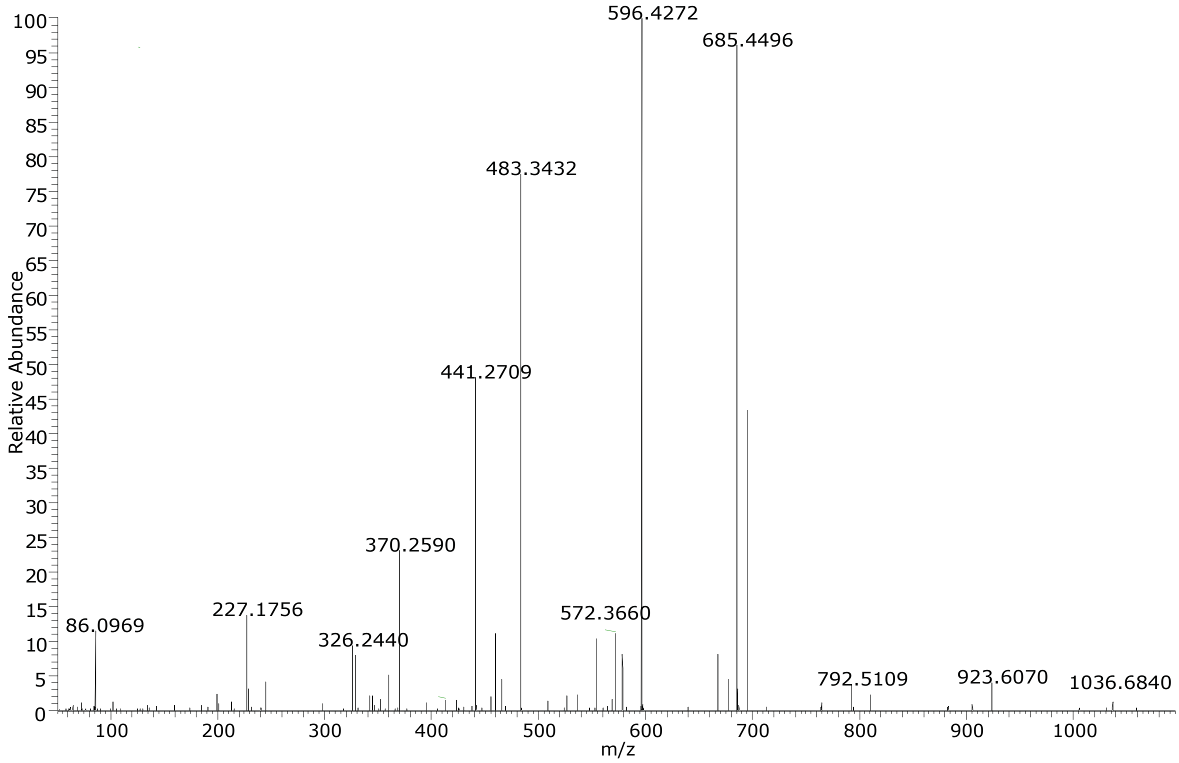

**Figure S 13** – MS/MS fragmentation pattern of $[M+H]^+$ 1054.7014 (gageostatin B) in positive mode.

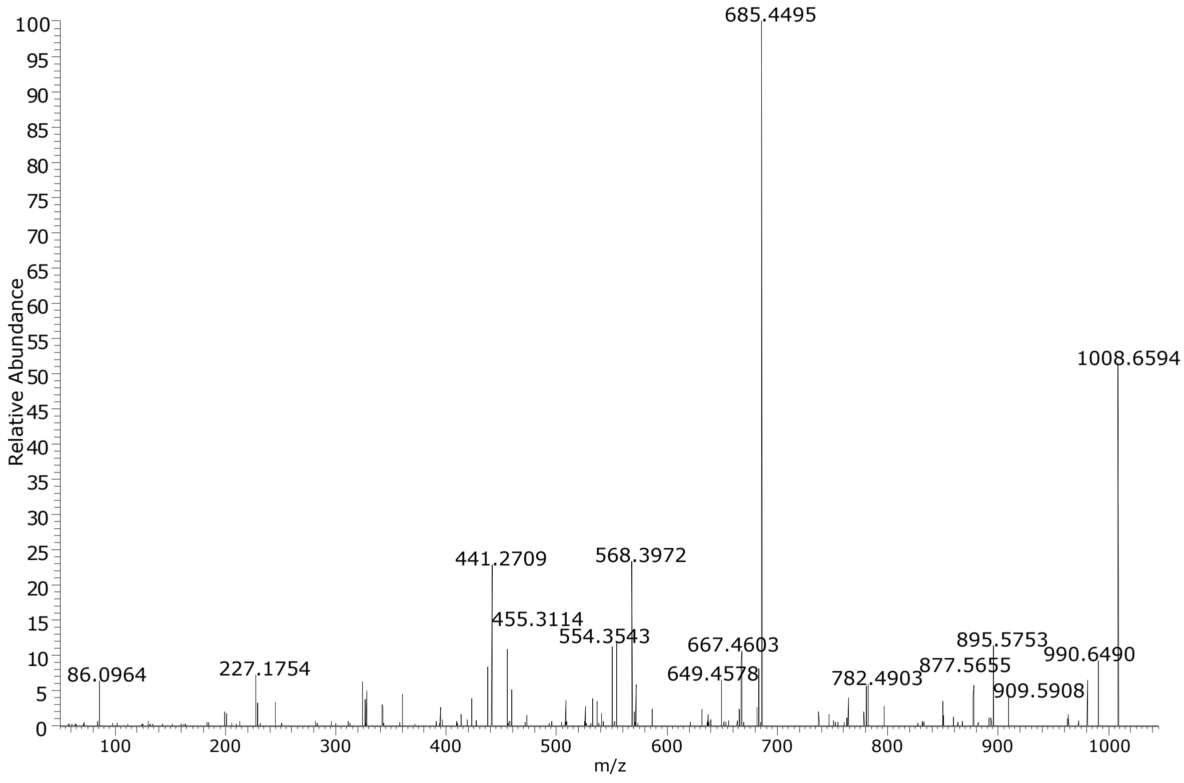

**Figure S 14** – MS/MS fragmentation pattern of $[M+H]^+$ 1008.6594 (gageostatin C) in positive mode.
